# scCapsNet: a deep learning classifier with the capability of interpretable feature extraction, applicable for single cell RNA data analysis

**DOI:** 10.1101/506642

**Authors:** Lifei Wang, Rui Nie, Ruyue Xin, Jiang Zhang, Jun Cai

## Abstract

Recently deep learning methods have been applied to process biological data and greatly pushed the development of the biological research forward. However, the interpretability of the deep learning methods still needs to improve. Here for the first time, we present scCapsNet, a totally interpretable deep learning model adapted from CapsNet. The scCapsNet model retains the capsule parts of CapsNet but replaces the part of convolutional neural networks with several parallel fully connected neural networks. We apply scCapsNet to scRNA-seq data. The results show that scCapsNet performs well as a classifier and also that the parallel fully connected neural networks function like feature extractors as we supposed. The scCapsNet model provides contribution of each extracted feature to the cell type recognition. Evidences show that some extracted features are nearly orthogonal to each other. After training, through analysis of the internal weights of each neural network connected inputs and primary capsule, and with the information about the contribution of each extracted feature to the cell type recognition, the scCapsNet model could relate gene sets from inputs to cell types. The specific gene set is responsible for the identification of its corresponding cell types but does not affect the recognition of other cell types by the model. Many well-studied cell type markers are in the gene set with corresponding cell type. The internal weights of neural network for those well-studied cell type markers are different for different primary capsules. The internal weights of neural network connected to a primary capsule could be viewed as an embedding for genes, convert genes to real value low dimensional vectors. Furthermore, we mix the RNA expression data of two cells with different cell types and then use the scCapsNet model trained with non-mixed data to predict the cell types in the mixed data. Our scCapsNet model could predict cell types in a cell mixture with high accuracy.

## Introduction

Single Cell RNA sequencing (scRNA-seq) could measure gene expression levels in individual cells. Using scRNA-seq data, it is possible to reveal heterogeneity in a cell population[1, 2], identify new cell types, computationally order cells along trajectories[3, 4], and infer the spatial coordinates of every individual cell in a population[5, 6].

As the scRNA-seq data accumulates quickly, it is important to retrieve similar cell types. For example, scMCA suggested a pipeline for cell type determination by comparing the input single-cell transcriptome with pre-calculated reference transcriptome to provide a match score based on gene expression correlation[7]. Since many cell types have already been well defined, supervised learning is an ideal tool to classify undefined cells. Besides the final goal of classification or similar cell type retrieval, the interpretability of the classification process is also important. By demonstrating which features are extracted for obtaining a specific decision and how these features contribute to the decision, the classifier could offer valuable information to the downstream operation such as biomarker discovery. Deep learning model is a proper tool to deal with vast and complex data such as RNA-seq data, but lacks of interpretability[8]. Therefore, there is a need for a model that utilizes the deep learning method with increased interpretability [9, 10].

Recent years, deep learning methods have been applied to process biological data [11–13]. Specifically, several deep learning models were used to analyze scRNA-seq data for various purposes. The neural networks which incorporate prior information were used to reduce the dimensions of the data[9] or discriminate tumor subtypes and their prognostic capability[10]. Variational inference (VI) method could achieve interpretable dimensionality reduction[14], single cell grouping[15], and approximating parameters which govern the distribution of expression value of each gene in each cell[16]. Generative adversarial network (GAN) was proved useful to simulate gene expression and predict perturbation in single cell[17].

The CapsNet is a novel deep learning model which is used in the task of digit recognition and exhibits more interpretability [18]. The CapsNet model could be used in protein structure classification and prediction [19, 20] and hold the great potential to apply in network biology and disease biology with data from multi-omics dataset [11].

Here for the first time, we propose a modified CapsNet model suitable for scRNA-seq data. In our experiment, we substitute the feature extraction part which uses convolutional neural networks in CapsNet with several parallel fully connected neural networks that use Rectified Linear Unit (ReLU) as the activation function. We reckon that, the parallel neural networks would function as a feature extractor. We call this modified model as “scCapsNet” and apply it to scRNA-seq data of mouse retinal bipolar cells [21] and human peripheral blood mononuclear cells (PBMC) [22]. We show that after training, the feature extraction part of scCapsNet could capture features from scRNA-seq data. And the capsule network part would compute the precise contribution of those captured features for cell type recognition. After training, through analysis of the internal weights of each neural network connected inputs and primary capsule, and with the information about the contribution of each extracted feature to the cell type recognition, the scCapsNet model could relate gene sets from inputs to cell types. The specific gene set is responsible for the identification of its corresponding cell types but does not affect the recognition of other cell types by the model. So genes in the gene set are vital for the cell type identity. We found that many well-studied cell type markers are among the gene set corresponding to the cell types. The internal weights of neural network for those well-studied cell type markers are different for different primary capsules, and give different kinds of representation. The internal weights of neural network connected to a primary capsule could be viewed as an embedding for genes, convert genes to real value low dimensional vectors, and analogize to the embedding of word (word2vec) in natural language processing [23]. With different requirements (primary capsules), the embedding is different. We also mix the expression data of two cells with different cell types together and use scCapsNet trained with non-mixed data to predict the types of the mixed data. The scCapsNet performs well to correctly predict the two cell types.

## Methods

### Datasets and data preprocessing

The two scRNA-seq datasets used in this work are mouse retinal bipolar neurons profiled by the Drop-Seq technology [21] and human peripheral blood mononuclear cells (PBMC) sequenced by 10X [22]. For data consistence, we directly adopt the data processing module from previous work[16]. After preprocessing, the mouse retinal dataset consists of 19829 cells with 13166 genes in each cell and the PBMC dataset consists of 11990 cells with 3346 genes in each cell. All expression data is converted into log-scale through log(x+1). We randomly divide the whole dataset into training set and validation set with ratio 9 to 1 by the method of train_test_split in the python package sklearn.model_selection.

### scCapsNet model

We adapt our scCapsNet model from previous CapsNet model. The architecture of our model is shown in Figure 1. The model contains two parts, one of which is feature detection part and the other is capsule network part. In the original CapsNet, the convolutional kernels were used as a local feature extractor for image input. Instead, we choose several parallel fully connected neural networks using Rectified Linear Unit (ReLU) activation function as the feature extractor. The output of each fully connected neural network is a vector with equal length of sixteen-dimension, and could be viewed as “primary capsule” in the original CapsNet. This part of our model converts the feature in RNA expression to the activities of local feature extractors. Next, the information would be delivered through primary capsule to the capsule in the final layer by “dynamic routing”. The capsule in the final layer， which corresponds to cell type and called “type capsule”, is a sixteen-dimension vector. The dynamic routing operation with three iterations occurs between the primary capsule layer and the type capsule layer. In our cell-type classification task, the length of the final layer type capsule represents the probability that one cell belongs to that cell type as the original CapsNet.

**Figure 1:**
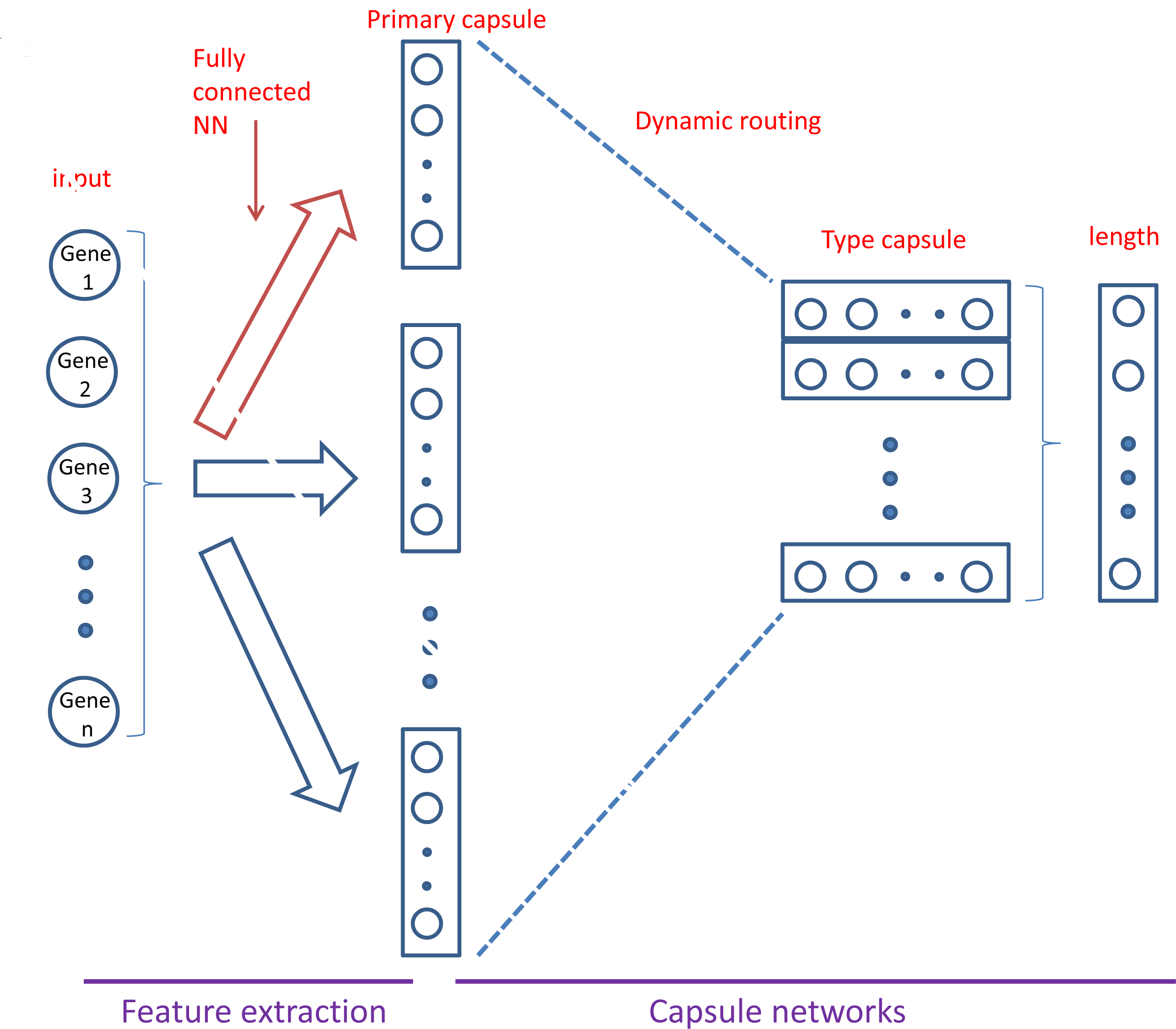
Architecture of scCapsNet with two layers. First layers consist of sixteen parallel fully connected neural networks for feature extraction. Each output of the neural network is viewed as a primary capsule. The subsequent layer is directly adopted from CapsNet for classification. Information flows from primary capsules to final layer type capsules. Then the length of each type capsule represents the probability of input data belonging to the corresponding cell type.

### Feature extraction and its contribution to type recognition

The detailed description for dynamic routing could be found in the original CapsNet paper[18]. In our scCapsNet model, the primary capsules that store the extracted features are first multiplied by weight matrices to produce “prediction vectors”. The dynamic routing process would calculate the “coupling coefficients” between each primary capsule’s prediction vectors and all type capsules. So the total number of coupling coefficients is the product of the number of primary capsules multiplied by the number of type capsules. The coupling coefficients of one primary capsule would sum up to one. After dynamic routing, the type capsule is actually a weighted sum of prediction vectors from primary capsules. The weights are the coupling coefficients and the magnitude of those coefficients indicates the contribution of the primary capsules to the type capsules. So the coupling coefficients could be views as the likelihood of the cell type containing the extracted feature stored in the primary capsules.

### Genes related with a specific feature and cell type

Between RNA-seq inputs and a primary capsule, a fully connected neural networks using Rectified Linear Unit (ReLU) activation function is used as a feature detector. After training, the internal weights of the neural networks are determinant. So, the weights associated with each gene (Figure S1) could be used as a label for each gene. Principle component analysis (PCA) is used to reduce the dimension of the internal weights of the neural networks. Then genes are plotted according to their first two principle components. The genes are chosen along one principle components. The effect of the exclusion of the chosen genes on cell type classification is measured by the prediction accuracy of each cell type.

### Neural networks for model comparison

In order to model comparison, we replace the capsule part in scCapsNet with fully connected neural networks. The concatenated feature extraction layer is directly connected to a fully connected neural network with final classification layer using sigmoid as activation function (**FigureS2**). The loss function is as the same as that of scCapsNet model.

### Mixed dataset and two-type detection

We mix the RNA expression of two different type cells with ratios of 1:1, 3:2, 2:1, 5:2 and 3:1. For type prediction of mixed data, all models are trained with non-mixed data. In order to make the models output two cell types, the top two largest scores in the output of both scCapNet and comparison model are selected, and only the correct prediction of both types is viewed as positive result. Each scalar between 0 and 1 in the output of both scCapNet and comparison model represents the probability that one cell belongs to one particular cell type. A threshold could be set for the second largest score, the cell which outputs a score below this threshold would be discarded. The prediction accuracy among the non-discarded cells would be calculated. Then the percentage that how many cells are not discarded is computed.

## Results

To test our model, two single cell RNA-seq datasets are used. One consists of mouse retinal bipolar neurons profiled by the Drop-Seq technology and the other consists of human peripheral blood mononuclear cells (PBMC) generated by 10X. Both datasets are randomly divided into training set and validation set with a ratio of 9:1. The model is run several times to access the prediction accuracy. For mouse retinal bipolar neurons dataset consisting of 19829 cells of 15 types and 13166 genes, the training accuracy reaches nearly 100% and the average validation accuracy is around 98%. For human PBMC dataset consisting of11990 cells of 9 types (we exclude cells with type “other”, so there were actually 8 types) and 3346 genes, the training accuracy reaches nearly 100% and the average validation accuracy is around96%.

### Primary capsule and cell type recognition

We want to find whether the primary capsule that output by feature extraction part could capture the properties in the single cell RNA-seq dataset. Furthermore, we want to explore how the extracted features contribute to the cell type recognition. The coupling coefficient could describe the relationship between the type capsule and the primary capsule (**method**). For our supervising learning task, we could group the coupling coefficients of the cells with prior known same cell type and calculate the average coupling coefficients for cells with one specific type. So we first compute the average coupling coefficients of each cell type in the validation set of the PBMC dataset and draw the heatmaps (Figure 2A). From the heatmaps, we find that for input cells with one specific cell type, one or several primary capsules contribute to the type recognition for this specific cell type but not obvious for other type. For example, among all the averaged coupling coefficients, the coupling coefficients of the primary capsule nine-B cells type capsule and the primary capsule fifteen-B cells type capsule are relatively higher than rest coupling coefficients for B cells input (Figure 2A**,0**). The coupling coefficient of the primary capsulezero-CD14+ monocyte type capsule is obviously higher than rest coupling coefficients (Figure 2A**, 1**).

**Figure 2A:**
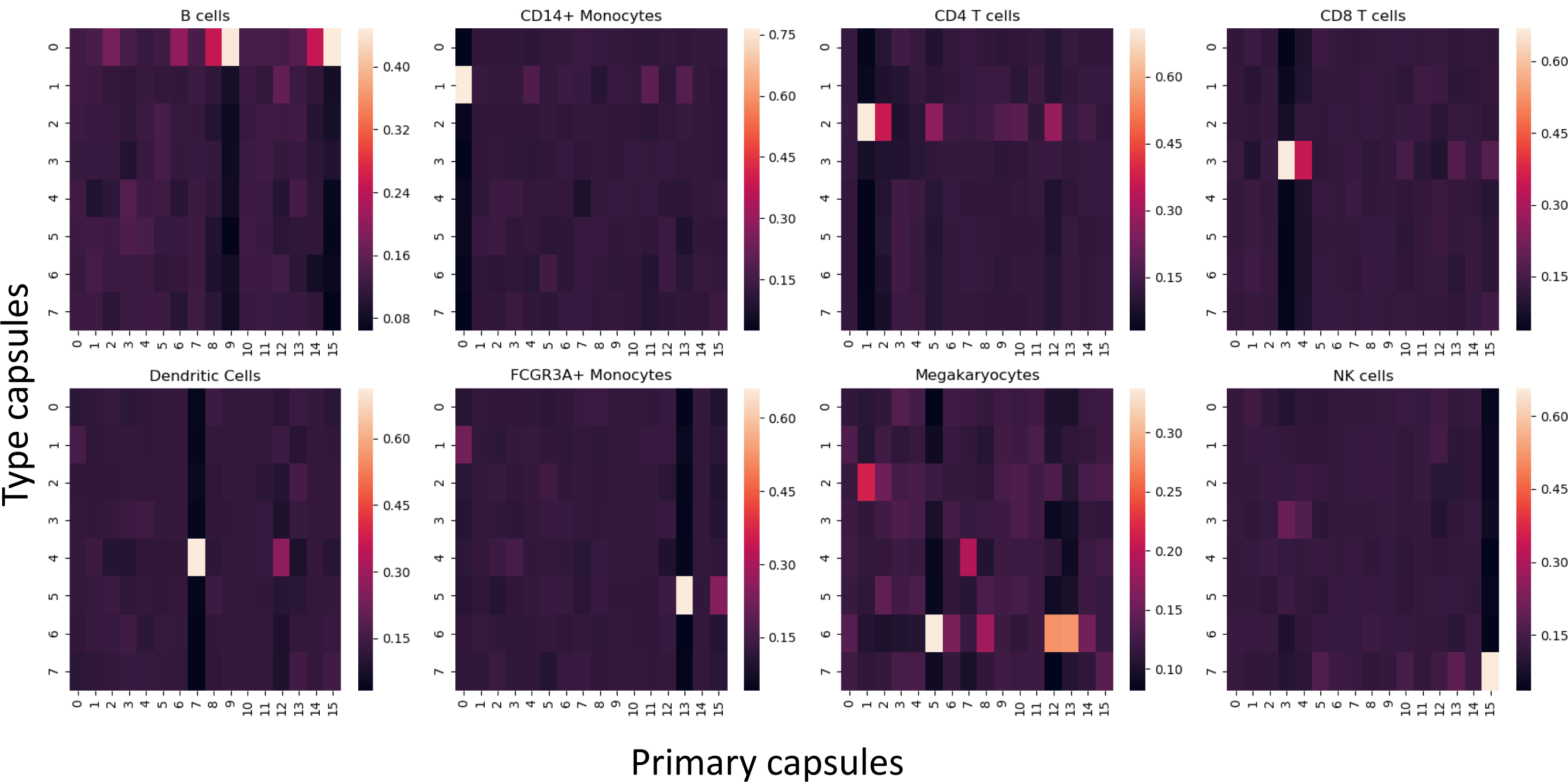
The heatmaps of coupling coefficients for PBMC dataset. Each heatmap represents the average coupling coefficients of inputs cells that belong to one specific cell type. In the heatmap, the row represents type capsules and column represents primary capsules. For example, the row zero represents B cells type capsule. The total number of the coupling coefficients is the product of the number of the type capsules multiplied by the number of the primary capsules. The order of the subplot is from left to right and top to bottom with index from zero to seven.

Next, in order to explore how one primary capsule affects the cell type recognition, we combine the average coupling coefficients with type capsules corresponding to specific cell types together into an overall heatmap (Figure 2B). We also use TSNE and PCA to reduce the dimension of each primary capsule, and plot the 2D-TSNE results (Figure 2C) and 2D-PCA results (Figure S3). We find some relationship between the overall heatmap of average coupling coefficients and the 2D-TSNE plot of each primary capsule. The details would be described hereafter.

**Figure 2B:**
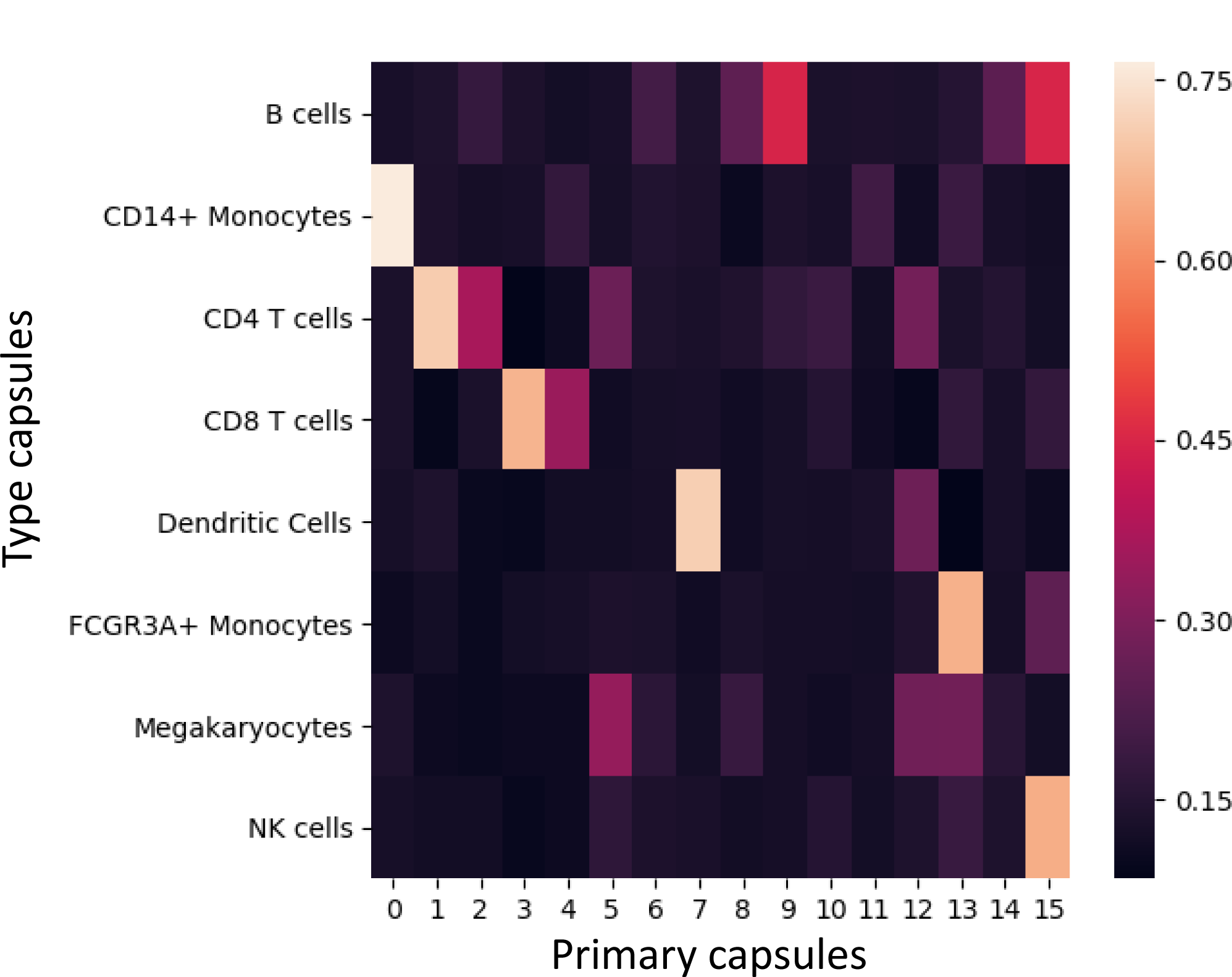
The overall heatmap of average coupling coefficients with type capsules corresponding to specific cell type inputs for PBMC dataset. The row represents type capsules and column represents primary capsules.

**Figure 2C:**
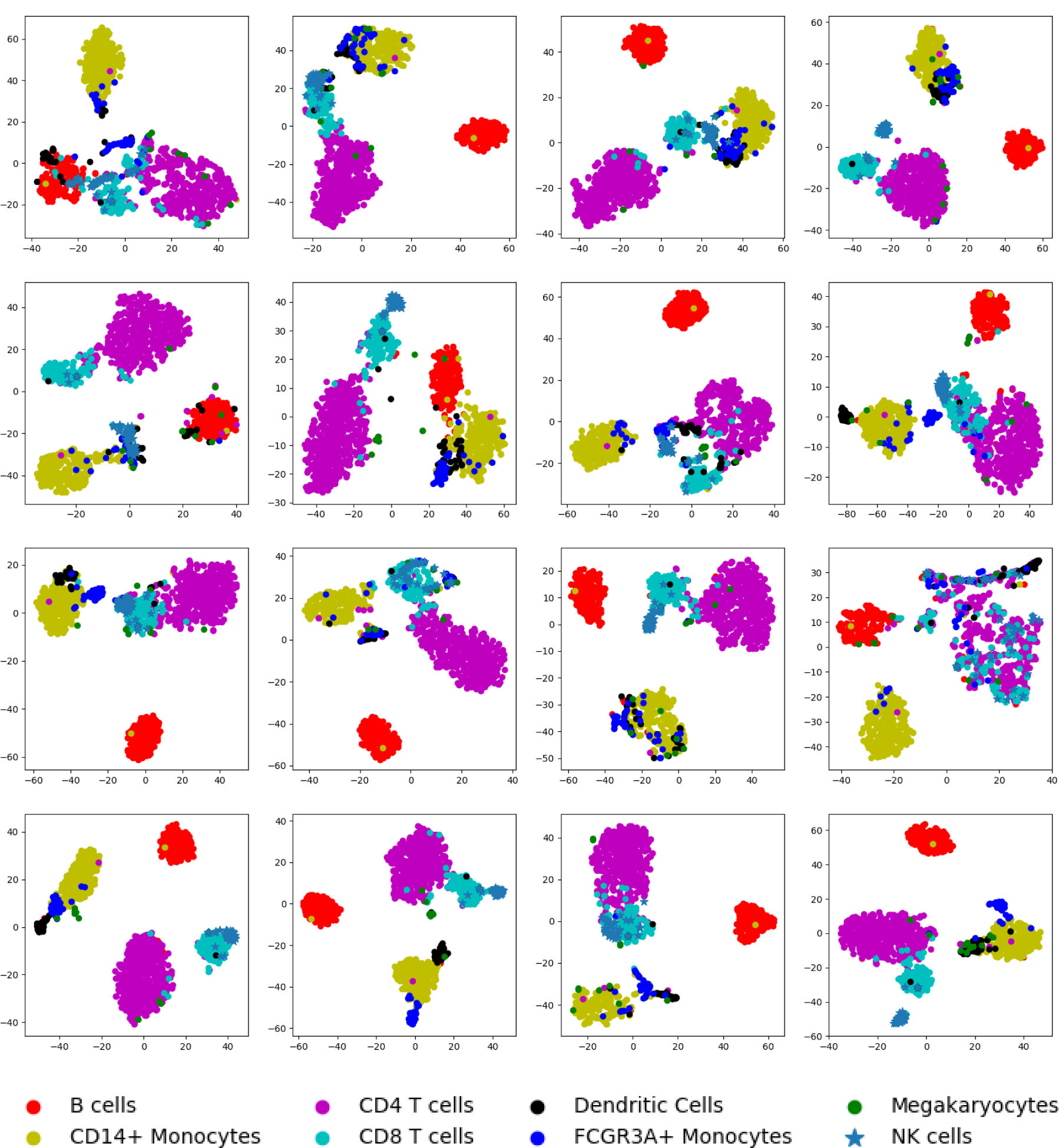
The 2D-TSNE plots of primary capsules of input cells from validation set for PBMC dataset. Each subplot represents a primary capsule. The order of the subplot is from left to right and top to bottom with index from zero to fifteen.

For example, the overall heatmap indicates the primary capsule zero contributes to the CD14+monocytes recognition, and the 2D-TSNE plot of primary capsule zero shows that the CD14+ monocytes are clustered together and locate on the top and cells of rest cell types are mixed together on the bottom (Figure 2C**, 0**). The overall heatmap indicates the primary capsule one and two contribute to the CD4 T cell recognition, and the 2D-TSNE plot of primary capsule one and two show that the CD4 T cells locate on the left bottom and are almost separate from the cells of other cell types (Figure 2C**, 1 and 2**). The overall heatmap indicates that the primary capsule three and four mainly contribute to the CD8 T cell recognition, and the 2D-TSNE plot of primary capsule three and four show that the CD8 T cells locate on the left and are separate from the cells of other cell types especially NK cells which normally could not be distinguished from CD8 T cells in most 2D-TSNE plot (Figure 2C, 3 and 4). The overall heatmap indicates that the primary capsule seven contributes to the dendritic cells recognition, and the 2D-TSNE plot of primary capsule seven shows that the Dendritic cells locate on the left and are almost separate from the cells of other cell types(Figure 2C**,7**). The overall heatmap indicates that the primary capsule eight and nine contribute to the B cells recognition, and the 2D-TSNE plot of primary capsule eight and nine show that the B cells locate on the bottom and are separate from the cells of rest cell types (Figure 2C**, 8 and 9**). The overall heatmap indicates that the primary capsule thirteen contributes to the FCGR3A+ monocytes recognition, and the 2D-TSNE plot of primary capsule thirteen shows that the FCGR3A+ monocytes locate on the bottom (Figure 2C**,13**). The overall heatmap indicates that the primary capsule fifteen contributes to the NK cells recognition, and the 2D-TSNE plot of primary capsule fifteen shows that the NK cells locate on the bottom and are separate from cells of other cell types especially CD8+ T cells (Figure 2C**,15**).

We also perform the same procedure on the mouse retina datasets. The heatmap of average coupling coefficients for input cells of prior known one specific cell type shows that, one or several primary capsules contribute to the type recognition for this specific cell type but not obvious for other type (Figure 3A). For example, among all the average coupling coefficients, the coupling coefficients of primary capsule fourteen-BC5A type capsule and primary capsule eleven-BC5A type capsule are obviously higher than other coupling coefficients for BC5A cells input (Figure 3A**,2**). The coupling coefficients of primary capsule five-BC7type capsule and primary capsule eleven-BC7 type capsule are obviously higher than other coupling coefficients for BC7 cells input (Figure 3A**,3**). The coupling coefficient of primary capsule ten-BC3A type capsule is obviously higher than other coupling coefficients for BC3A cells input (Figure 3A**, 11**). The coupling coefficient of primary capsule fourteen-BC5D type capsule is obviously higher than other coupling coefficients for BC5D cells input (Figure 3A**, 10**).

**Figure 3A:**
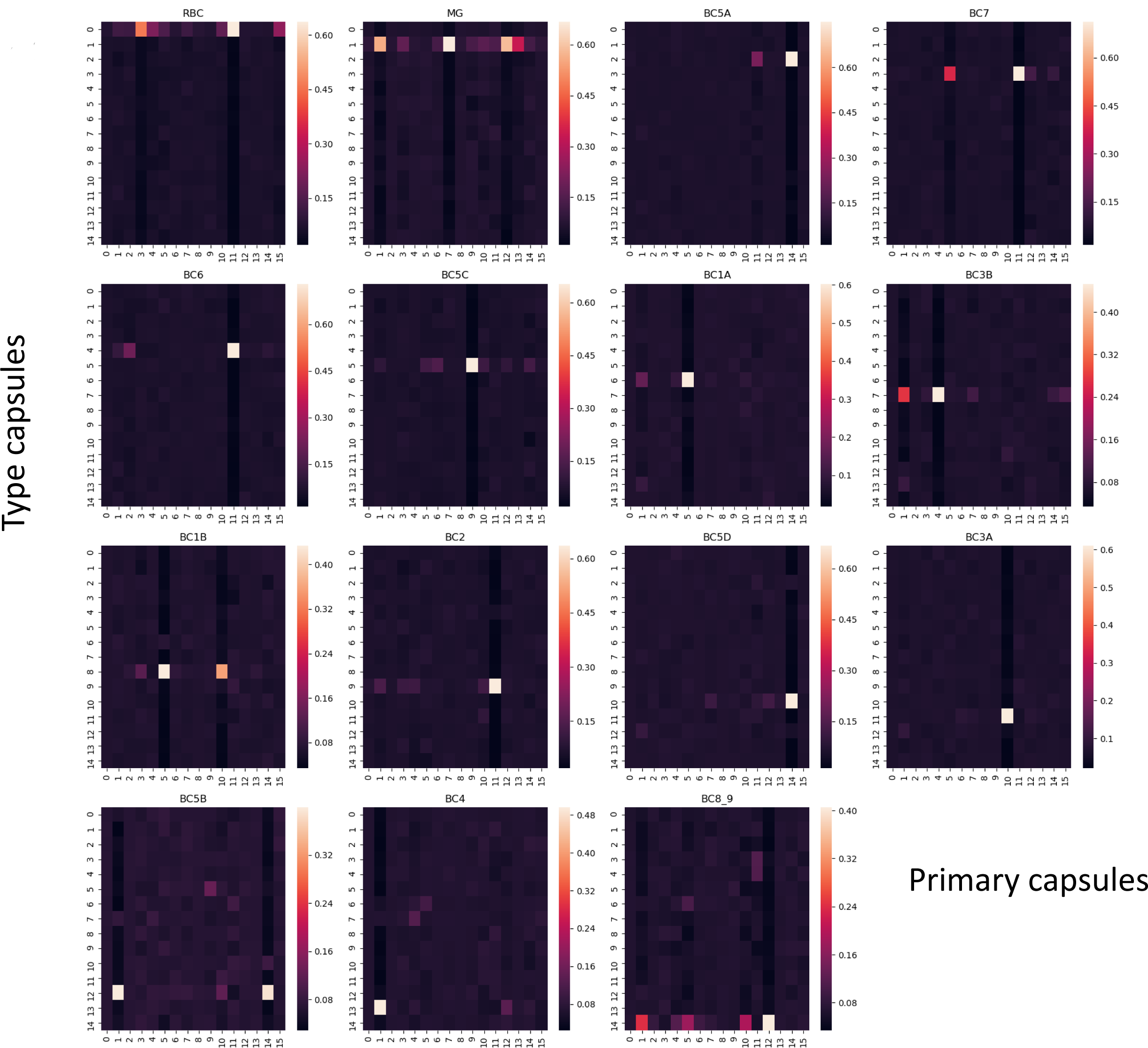
The heatmaps of coupling coefficients for mouse retina bipolar cells dataset. Each heatmap represents average coupling coefficients of inputs cells that belong to one specific cell type. In the heatmap, the row represents type capsules and column represents primary capsules. For example, the row zero represents the RBC type capsule. The total number of the coupling coefficients is the product of the number of the type capsules multiplied by the number of the primary capsules. The order of the subplot is from left to right and top to bottom with index from zero to fourteen.

We also combine the average coupling coefficient with type capsules corresponding to specific cell type inputs together into an overall heatmap (Figure 3B) and draw the 2D-TSNEand 2D-PCA plot of outputs of every primary capsule (Figure 3C **and** Figure S4). The overall heatmap indicates that the primary capsule two contributes to the BC6 cell recognition, and the 2D-TSNE plot of primary capsule two shows that most BC6 cells are clustered together on the top and cells of other cell types especially cone cells are mixed together on the bottom (Figure 3C**,2**). The overall heatmap indicates that the primary nine contributes to the BC5C cell recognition, and the 2D-TSNE plot of primary capsule nine shows that the BC5C cells are clustered on the top and cells of other cell types especially cone cells are mixed together on the bottom (Figure 3C**, 9**). The overall heatmap indicates that the primary four contributes to the BC3B and RBC cell recognition, and the 2D-TSNE plot of primary capsule four shows that the BC3B and RBC cells locate on the left and are almost separate from the rest cells (Figure 3C**,4**). The overall heatmap indicates that the primary five contributes to the BC1A, BC1B and BC7 cell recognition, and the 2D-TSNE plot of primary capsule five shows that the BC1A, BC1B and BC7 cells located on the right and are almost separate from the rest cells (Figure 3C**, 5**). The overall heatmap indicates that the primary capsule ten contributes to the BC1B and BC3A cell recognition, and the 2D-TSNE plot of primary capsule ten shows that the BC1B and BC3A cell locate on the bottom and are almost separate from the rest cells (Figure 3C**, 10**). The overall heatmap indicates that the primary fourteen contributes to the BC5D, BC5B and BC5A cell recognition, and the 2D-TSNE plot of primary capsule fourteen shows that the BC5D, BC5B and BC5A cells locate on the left and are almost separate from the rest cells (Figure 3C**, 14**). The heatmap indicates that the primary capsule eleven contributes to the BC2, BC6, BC7, BC5A and RBC cell recognition, and the 2D-TSNE plot of primary capsule eleven shows that the BC2, BC6, BC7, BC5A and RBC cells are independently clustered for their own cell type and cells of other cell types are mixed together (Figure 3C**, 11**).

**Figure 3B:**
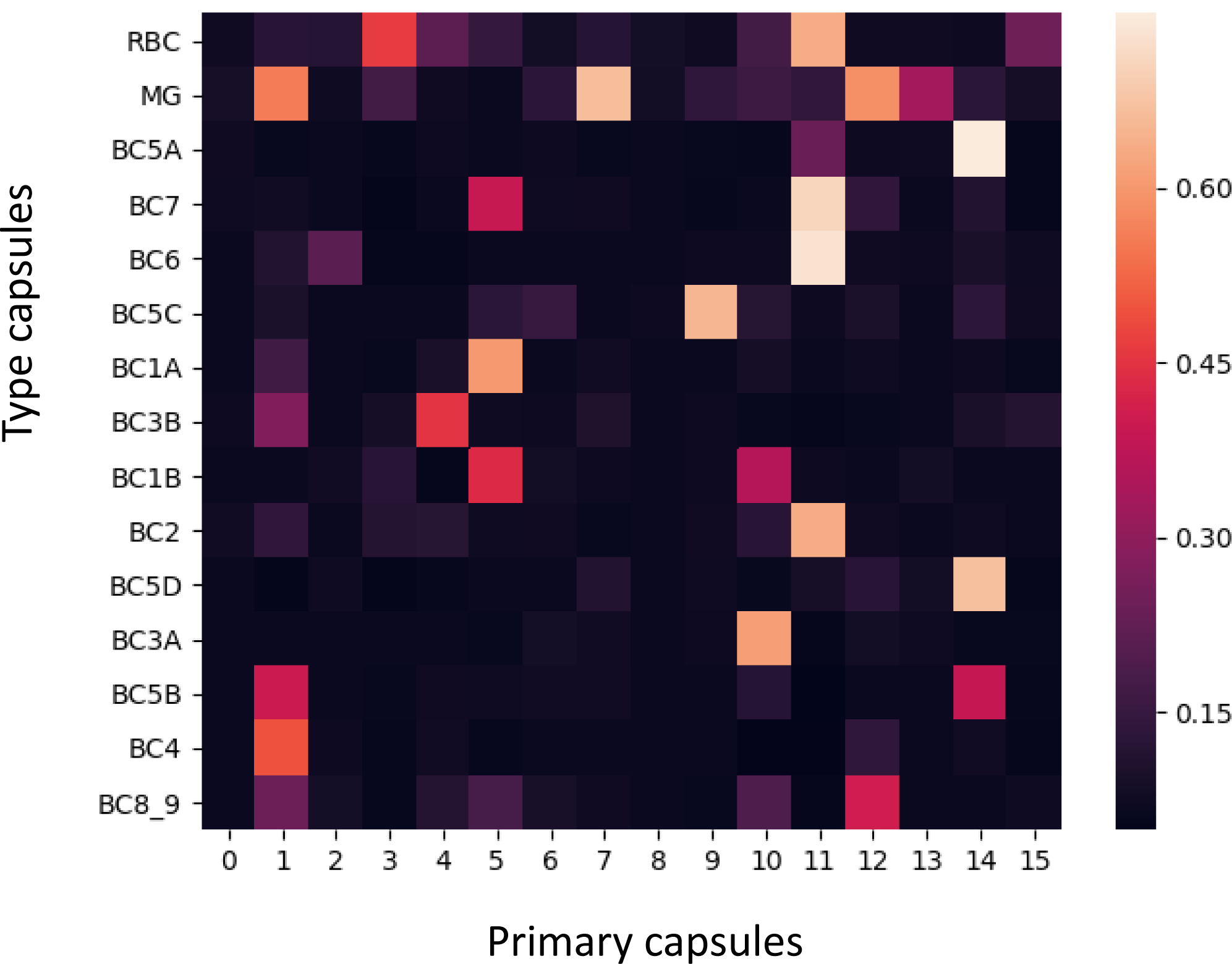
The overall heatmap of average coupling coefficients with type capsules corresponding to a specific cell type input for mouse retina bipolar cells dataset. The row represents type capsules and column represents primary capsules.

**Figure 3C:**
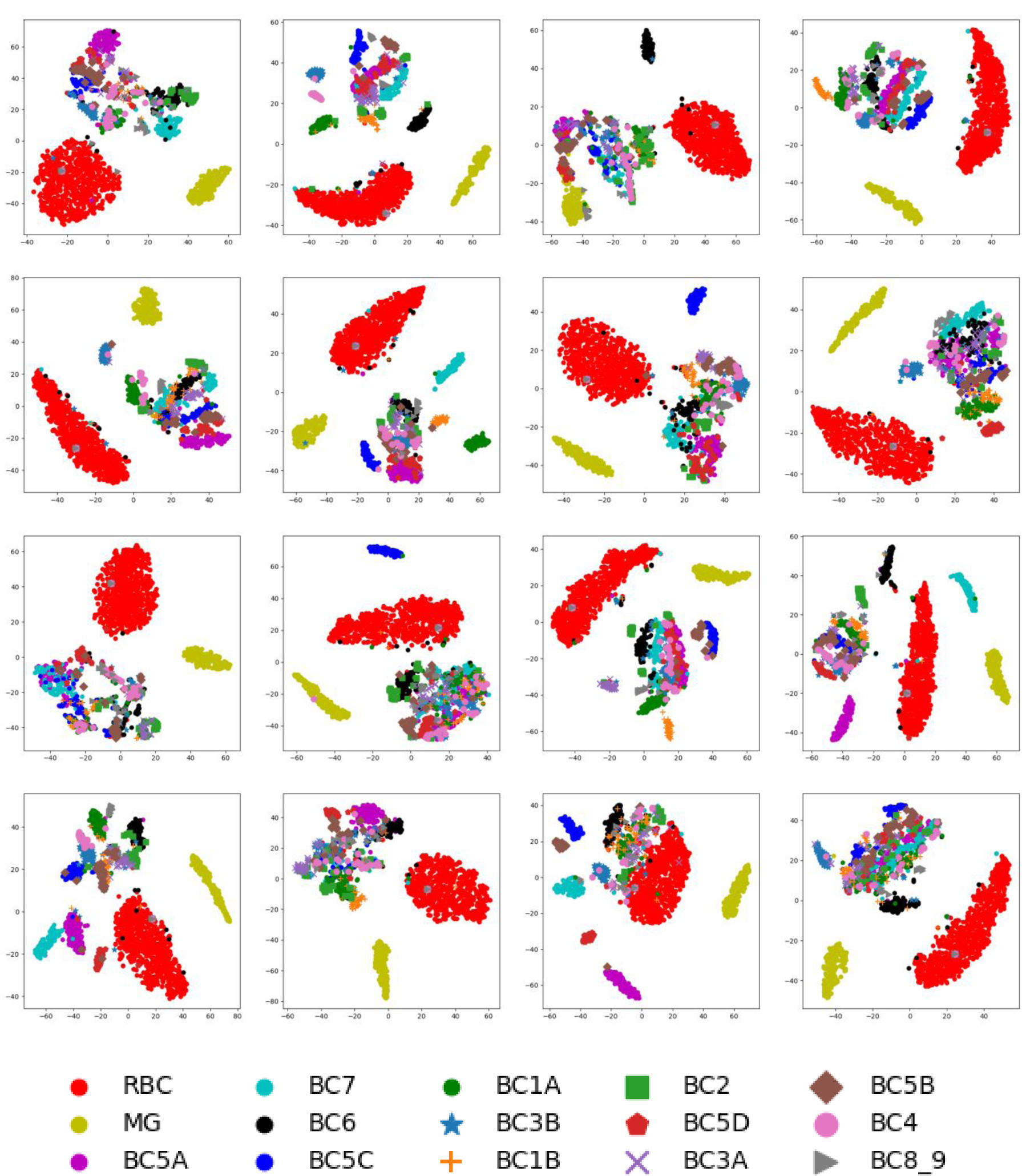
The 2D-TSNE plots of primary capsules of input cells from validation set for mouse retina bipolar cells dataset. Each subplot represents a primary capsule. The order of the subplot is from left to right and top to bottom with index from zero to fifteen.

From the above observation, we find the association between heatmap of average coupling coefficients and 2D-TSNE plot of primary capsules. If the heatmap indicates one primary capsule contributes to certain cell type recognition, the cells belong to those cell types would independently cluster into separate groups, while cells labeled as other cell types would mix together. This phenomenon reveals that the primary capsule actually captures some properties of scRNA-seq datasets, and contribution of those captured properties to the cell type recognition could be measured by coupling coefficients.

### Specific gene set, primary capsule and cell type recognition

After model training, the internal weights of the neural network (Figure S1) that connect inputs and primary capsule are determinant. For a primary capsule, the internal weights associated with a gene could be views as a label for that gene. Furthermore, this real-value low dimension label actually represents an embedding which embeds the gene into low dimension space according to the needs for a specific primary capsule. Then, we perform PCA on internal weights of the neural network for a primary capsule and choose genes along one principle component in order to find their role in the cell type recognition.

We use primary capsule three, which is related to CD8 T cell recognition (Figure 2B), for demonstration. As show in Figure 4A, the genes are plotted according to their first two principle components. We choose genes along the principle component one, and set the inputs value of the chosen genes to zeros. Then this deficit dataset, with inputs value of many genes being set to zero, and the untouched dataset are both fed into trained model. We found that when setting the inputs value of several genes to zero (Figure 4A**, blue dots**), the trained model almost could not recognize the CD8 T cell while the ability to identify other cell types is almost not affected (Figure 4D). Furthermore, we also draw heatmaps of average coupling coefficients (Figure 4B) for deficit dataset and combine the average coupling coefficients with type capsules corresponding to specific cell types together into overall heatmap (Figure 4C). The heatmaps and the overall heatmap show that except CD8 T cells, other cells are not affected by the exclusion of several genes (setting gene value to zero), comparing to untouched dataset (Figure 2A, 2B). For CD8 T cells, the exclusion of several genes in the inputs prevents them from being identified as CD8 T cells. Instead, CD8 T cells are recognized CD4 T cells and natural killer cells (Figure 4E), due to the similarity of remaining genes. The heatmaps shows that the CD8 T cells inputs without specific genes could displace the characteristic of either CD4 T cells or NK cells (Figure 4B**, 3**), although with much lower scores (Figures 4B, 2 and 7).

**Figure 4:**
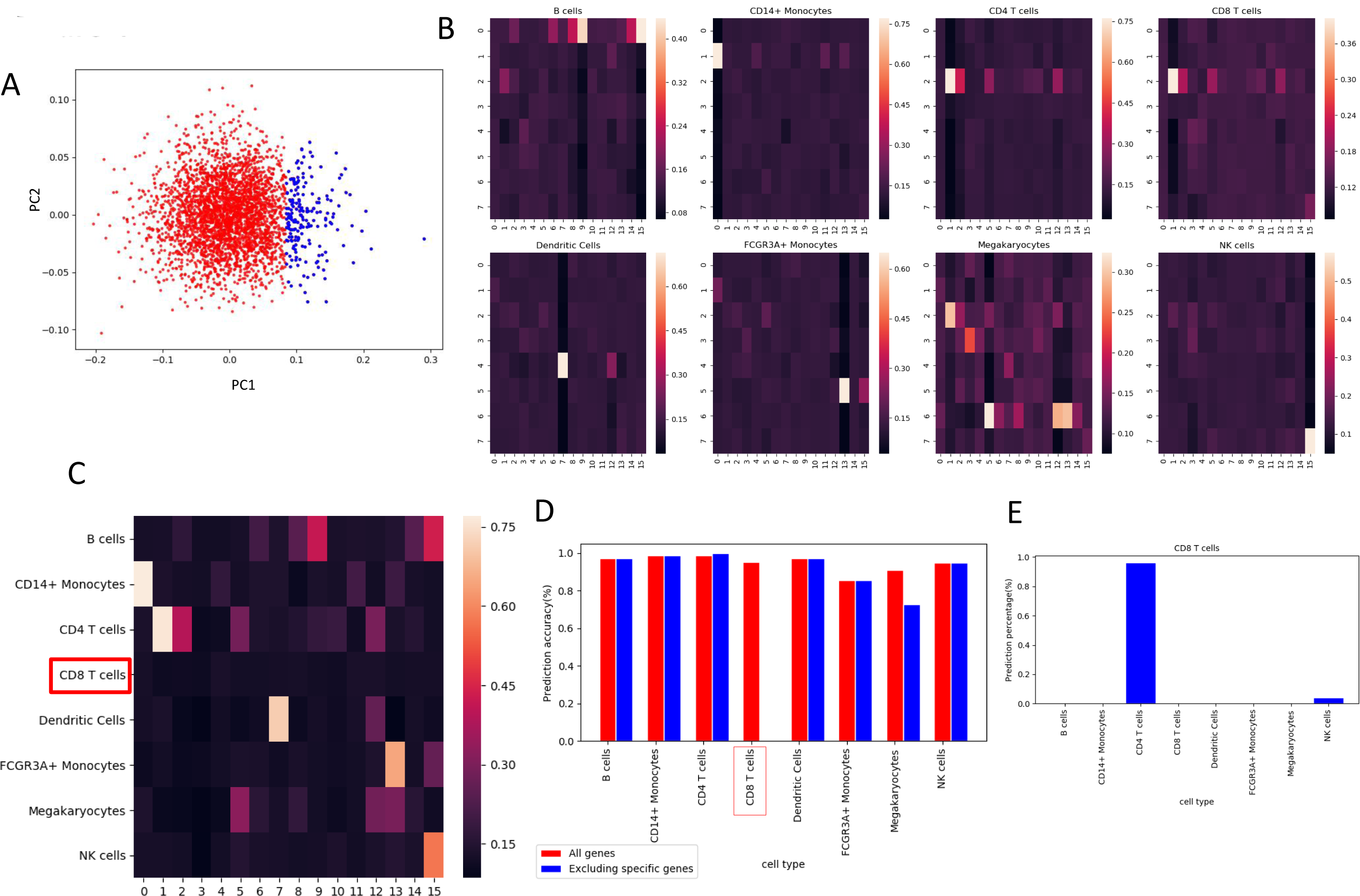
The identification of the gene set specific for CD8 T cell recognition. **A**. The plot of the first two principle components (PC) of PCA which perform on internal weights of neural network that connect inputs and primary capsule three. Each dot represents a gene. Blue dots are chosen for genes exclusion. **B**. The heatmaps of coupling coefficients for deficit PBMC dataset in which the inputs value of blue dot genes are set to zero. **C**. The overall heatmap of average coupling coefficients with type capsules corresponding to a specific cell type input for deficit PBMC dataset. **D**. The comparison of prediction accuracy of each cell type between the untouched PBMC dataset and the deficit PBMC dataset. **E**. The proportion misclassified cell types of CD8 T cells with deficit PBMC dataset.

All evidence above shows that the set of specific genes (Figure 4A**, blue dots**) are vital for CD8 T cells identity. The exclusion of those genes prevents model to correctly recognize CD8 T cells, while barely affects the recognition of cells with other cell types. With the same procedure, we could identify the gene set specific for each cell type in PBMC dataset (**Figures S5-S10,** Figure 5). One specific gene set is vital to one cell type recognition and almost does not affects the recognition of other cell types (**Figures S5-S10 and** Figure 5**, D**). With the loss of the essential genes for their identity, most B cells are misclassified as CD4 T cells (Figure S5**, F**), most CD4 T cells are identified as CD8 T cells (Figure S7**, F**), most DC cells are identified as FCGR3A+ monocytes (Figure S8**, F**), most FCGR3A+ monocytes are identified as CD14+ monocytes (Figure S9**, F**), most Natural killer cells are identified as CD8T cells (Figure 5**, F**).

**Figure 5:**
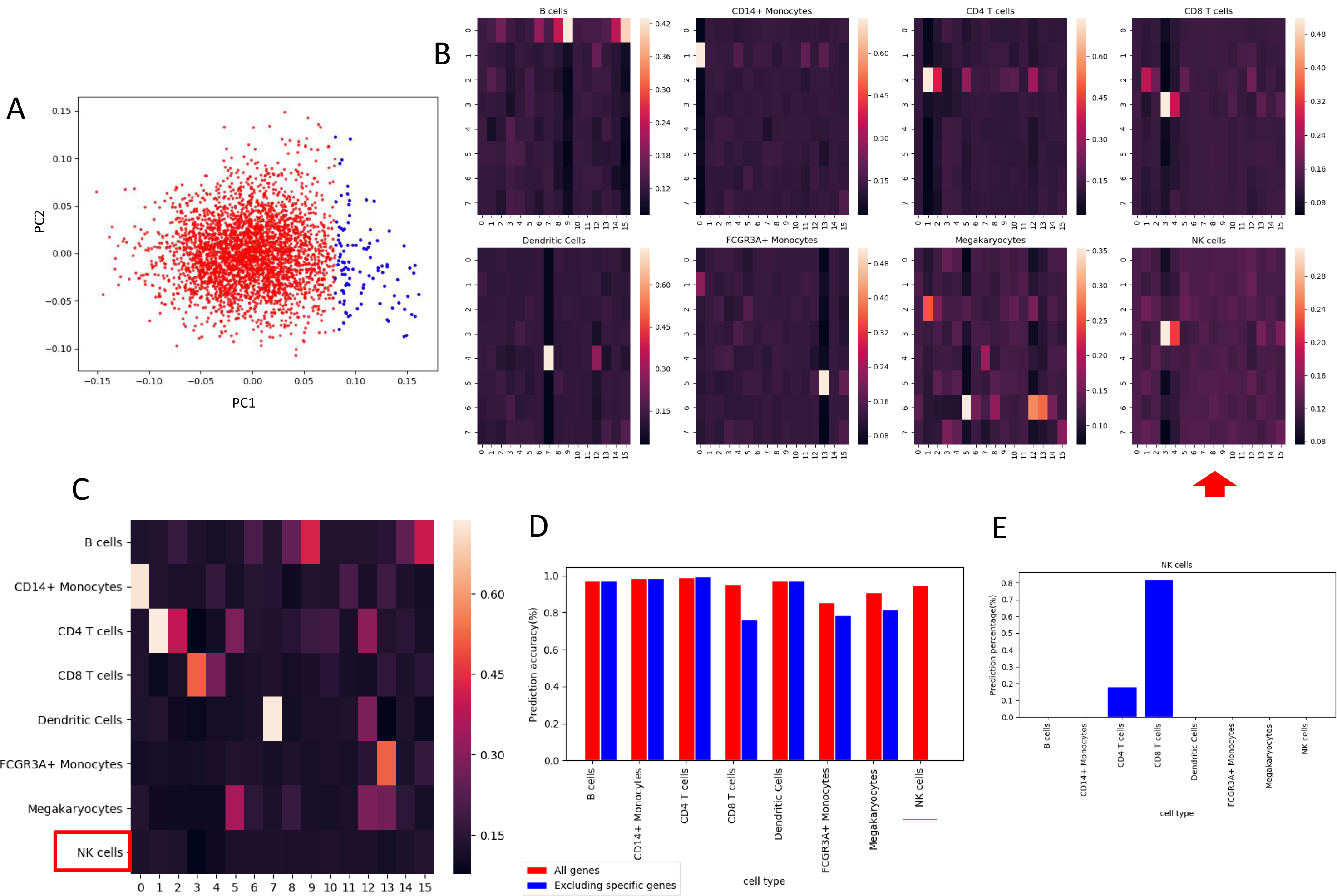
The identification of the gene set specific for NK cell recognition. **A**. The plot of the first two principle components (PC) of PCA which perform on internal weights of neural network that connect inputs and primary capsule fifteen. Each dot represents a gene. Blue dots are chosen for genes exclusion. **B**. The heatmaps of coupling coefficients for deficit PBMC dataset in which the inputs value of blue dot genes are set to zero. **C**. The overall heatmap of average coupling coefficients with type capsules corresponding to a specific cell type input for deficit PBMC dataset. **D**. The comparison of prediction accuracy of each cell type between the untouched PBMC dataset and the deficit PBMC dataset. **E**. The proportion misclassified cell types of NK cells with deficit PBMC dataset.

We also found similar phenomenon on mouse retinal bipolar cells dataset (Figure S11). A specific gene set is identified that associates with BC3B cells (Figure S11**, A**). With the exclusion of genes from this gene set, the ability of the trained model to recognize BC3B cells is significant damage, while other functions of the trained model are still largely retained (Figure S11**, B-E**).

The primary capsule fifteen seems at least responsible for the recognition of Natural killer cells and B cells (Figure 2B). The exclusion of several genes along principle component one axis (Figure 5**, A, blue dots**) prevent trained model recognizing Natural killer cells (Figures 5, C and D) and misclassify them as CD8 T cells (Figures 5, B and E). Furthermore, the exclusion of several genes along principle component two axis (Figure 6**, A, blue dots**) prevents trained model recognizing B cells (Figure 6, C and D) and misclassifies most of them as CD4 T cells (Figure 6, B and E). The example of primary capsule fifteen demonstrates that one primary capsule could be responsible for multiple type recognition by store different features in different orientation and different features may be orthogonal to each other.

**Figure 6:**
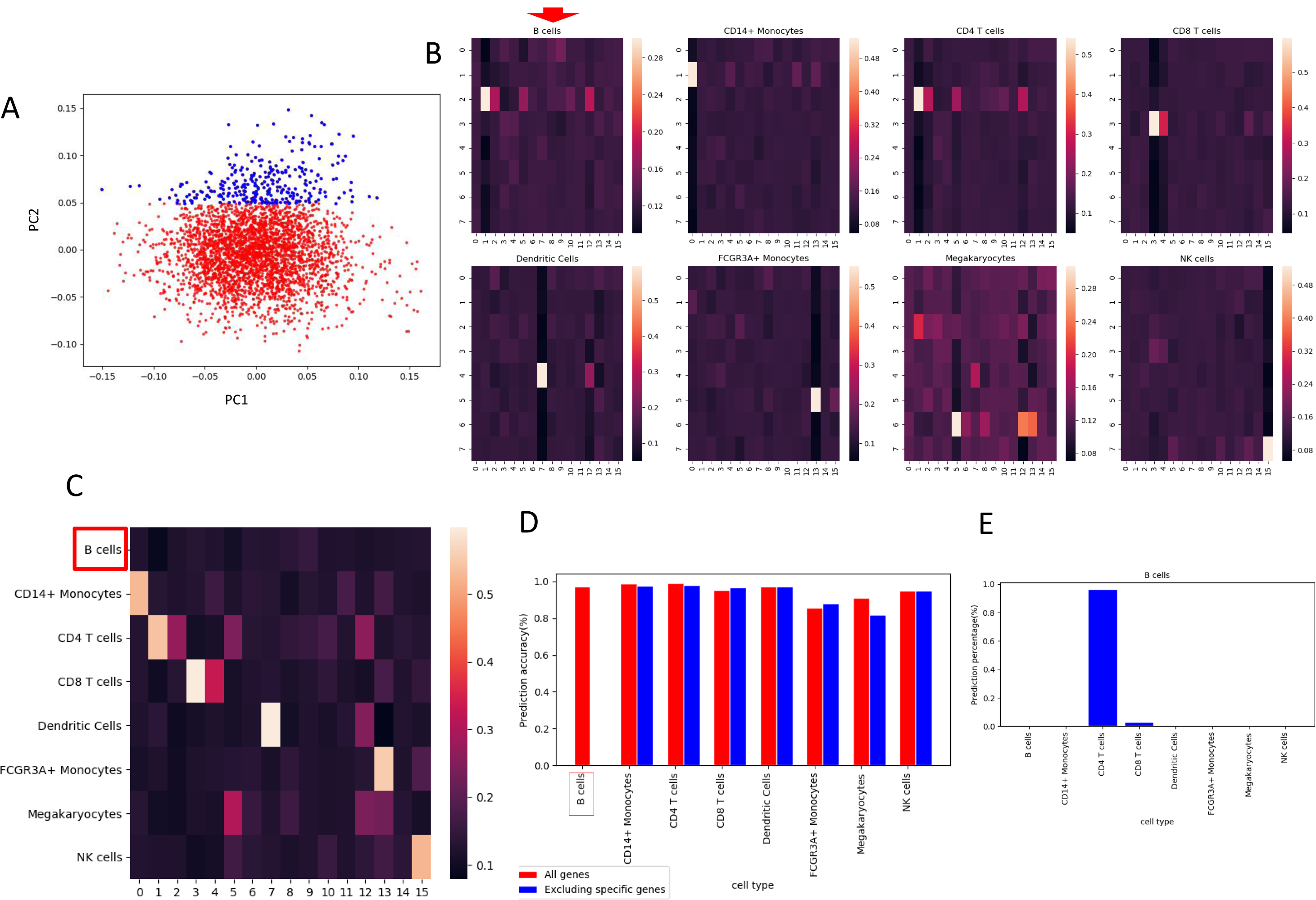
The identification of the gene set specific for B cell recognition. **A**. The plot of the first two principle components (PC) of PCA which perform on internal weights of neural network that connect inputs and primary capsule fifteen. Each dot represents a gene. Blue dots are chosen for genes exclusion. **B**. The heatmaps of coupling coefficients for deficit PBMC dataset in which the inputs value of blue dot genes are set to zero. **C**. The overall heatmap of average coupling coefficients with type capsules corresponding to a specific cell type input for deficit PBMC dataset. **D**. The comparison of prediction accuracy of each cell type between the untouched PBMC dataset and the deficit PBMC dataset. **E**. The proportion misclassified cell types of B cells with deficit PBMC dataset.

### Decomposition of features, embedding of genes into low dimensional space

Since several gene sets and their associated cell types have been identified, the question emerged immediately is that how the well-studied cell type markers are represented in trained scCapsNet model. We choose CD8A for CD8 T cells, CD14 for CD14+ monocytes and CD19 for B cells as examples for demonstration. We plot the first two principle components of PCA for internal weights of neural network connecting to primary capsule three (CD8 T cells), primary capsule one (CD14 monocytes) and primary capsule nine (B cells), then mark the positions of CD8A, CD14 and CD19 with colored stars on the plots (Figure 7). The plots show that each well-studied cell type maker is included in the specific gene set for their corresponding cell type identified by model, while is not in the specific gene set for other cell types. The embedding of those maker genes is different for different primary capsule.

**Figure 7:**
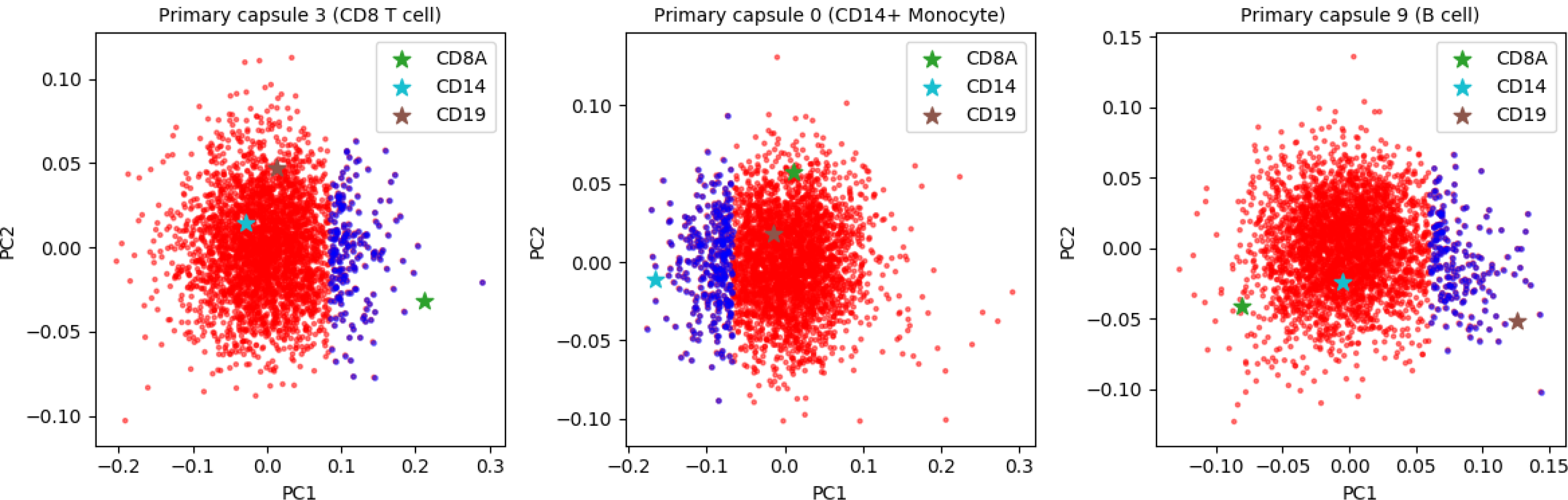
The positions of three well-studied cell type marker on the plot of principle components of PCA which preform on internal weights of neural network that connect inputs and primary capsule.

In order to get a big picture about how the genes are embedded for different primary capsules. We combine the weight of neural networks of primary capsule zero, one, three, five, seven, nine, thirteen and fifteen (Figure 2B), which we investigate before (Figures 4, 5 and S5-S10), together and perform PCA. The explained variance and explained variance ratio for each principle component is drawn (Figure 8A). Then we plot the different pair of principle component for genes with color and shape corresponding to the primary capsules, and name each primary capsule with its most responsible cell type (Figure 8B). The plots show that the embedding of genes from a particular primary capsule could be well explained by a principle component. For example, the embedding of genes from primary capsule one (CD4 T cell) and principle component two (Figures 8B, **top left, bottom left and bottom middle**) are highly correlated. And the embedding of genes from primary capsule fifteen (CD4 T cell) and principle component four (Figure 8B**, top left, bottom left and bottom middle**) are highly correlated. Those relationships indicate that the some features extracted by scCapsNet model are nearly orthogonal to each other.

**Figure 8:**
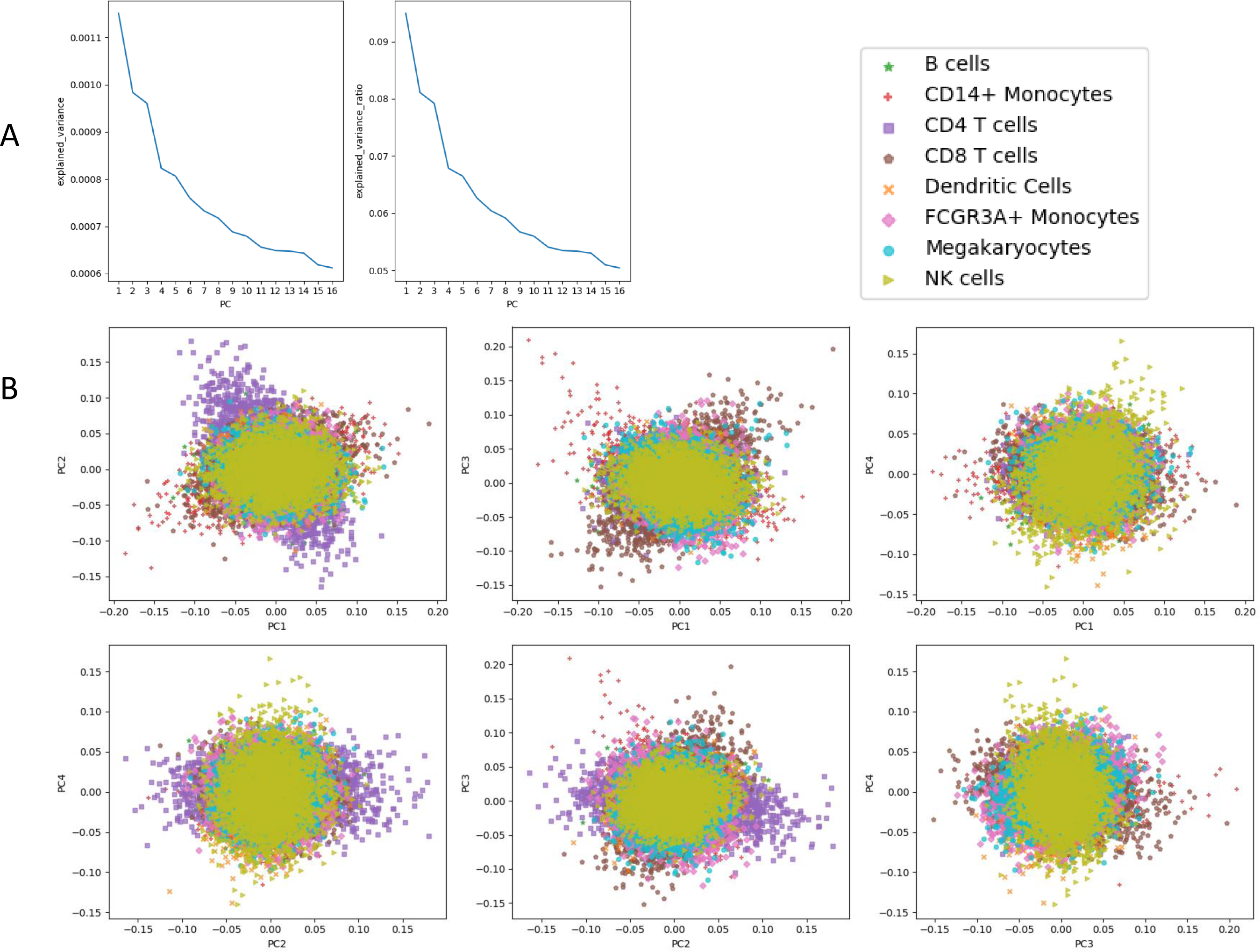
The analysis of embedding for genes associated with different primary capsules through PCA. **A**. The explained variance and explained variance ratio for each principle component. **B**. The plots of different pairs of principle components. The labels of each dot are associated with the corresponding primary capsules which named by their related cell types.

We then mark the CD8A (Figure S12), CD14 (Figure S13) and CD19 (Figure S14) on the plot of combined weights (Figure 8B). The position of each gene are scattered around the whole plot and indicate that each gene is embedded differently from different primary capsule. So each primary capsule assigns a particular embedding strategy that compel genes embed in a low dimension space, in order to fulfill their role in the cell type classification process.

### Mixture recognition

In hinton’s paper, the dynamic routing made the CapsNet capable of recognizing multiple objects in an image even if the objects were overlapped, although the model was trained with the single non-overlapped objects in image[18]. We mimic the overlapped objects in an image by adding the gene expression of two cells with different ratio in single cell RNA-seq data. We want to test whether our model trained with non-mixed data could decouple the data which is a mixture of two different types of cells. The results show that our model could accomplish the cell type prediction of the mixture with high accuracy. As we expect, the more unequal the cell mixture is, the lower prediction accuracy the model could make (Table 1, 2).

Although the overall prediction accuracies of our scCapsNets model are slightly lower than those of the comparison model, when we set a threshold to the results, we could easily find that the comparison model tends to give a lower probability as compared to scCapsNet. Under the same threshold, the percentage that the cells which output a score above this threshold by scCapsNet are obviously larger than the corresponding percentage of the comparison model (Table 1, 2).

**Table 1:** The performance of different models on the task of cell mixture types recognition for PBMC dataset. The threshold is set for the second largest score in the output of the model. The scCapsNet is our model, and NN is the comparison model which uses the fully connected neural networks to replace the capsule networks. The percentage is calculated by the number of the cells those are not filtered out by the threshold divided by the total number of cells. The accuracy is calculated among the cells that are not filtered out by the threshold.

|  | threshold | 0.5 |  | 0.4 |  | 0.3 |  | 0.2 |  | 0.1 |  |
| --- | --- | --- | --- | --- | --- | --- | --- | --- | --- | --- | --- |
| mixture | % | scCaps<br>Net | NN | scCaps<br>Net | NN | scCaps<br>Net | NN | scCapsN<br>et | NN | scCapsN<br>et | NN |
| 1:1 | Accuracy | 0.992 | 0.993 | 0.989 | 0.986 | 0.978 | 0.98 | 0.955 | 0.959 | 0.927 | 0.925 |
|  | percentage | 0.157 | 0.073 | 0.3 | 0.195 | 0.469 | 0.364 | 0.658 | 0.593 | 0.852 | 0.829 |
| 3:2 | Accuracy | 0.983 | 0.974 | 0.975 | 0.982 | 0.962 | 0.968 | 0.942 | 0.949 | 0.905 | 0.908 |
|  | percentage | 0.113 | 0.056 | 0.215 | 0.145 | 0.377 | 0.285 | 0.565 | 0.492 | 0.795 | 0.774 |
| 2:1 | Accuracy | 0.959 | 0.961 | 0.954 | 0.959 | 0.93 | 0.953 | 0.912 | 0.919 | 0.874 | 0.878 |
|  | percentage | 0.058 | 0.028 | 0.139 | 0.091 | 0.254 | 0.196 | 0.434 | 0.371 | 0.687 | 0.675 |
| 5:2 | Accuracy | 0.907 | 0.912 | 0.907 | 0.932 | 0.888 | 0.919 | 0.87 | 0.877 | 0.835 | 0.842 |
|  | percentage | 0.038 | 0.016 | 0.085 | 0.056 | 0.178 | 0.139 | 0.328 | 0.286 | 0.592 | 0.577 |
| 3:1 | Accuracy | 0.841 | 0.892 | 0.838 | 0.879 | 0.819 | 0.867 | 0.831 | 0.831 | 0.788 | 0.799 |
|  | percentage | 0.027 | 0.013 | 0.06 | 0.039 | 0.127 | 0.1 | 0.256 | 0.233 | 0.524 | 0.509 |

**Table 2:**
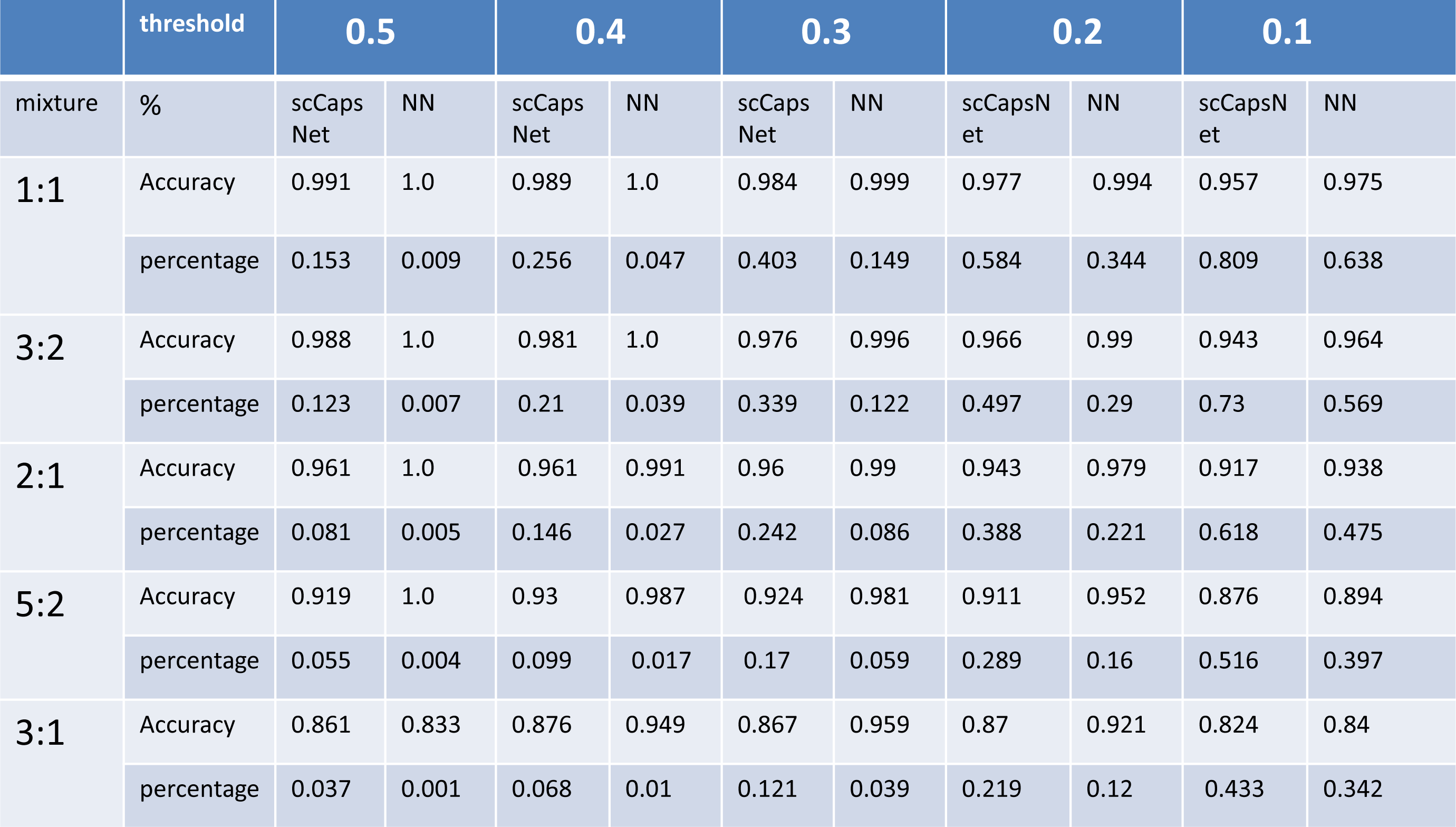
The performance of different models on the task of cell mixture types recognition for mouse retina bipolar cell dataset. The threshold is set for the second largest score in the output of the model. The scCapsNet is our model, and NN is the comparison model which uses the fully connected neural networks to replace the capsule networks. The percentage is calculated by the number of the cells those are not filtered out by the threshold divided by the total number of cells. The accuracy is calculated among the cells that were not filtered out by the threshold.

|  | threshold | 0.5 |  | 0.4 |  | 0.3 |  | 0.2 |  | 0.1 |  |
| --- | --- | --- | --- | --- | --- | --- | --- | --- | --- | --- | --- |
| mixture | % | scCaps<br>Net | NN | scCaps<br>Net | NN | scCaps<br>Net | NN | scCapsN<br>et | NN | scCapsN<br>et | NN |
| 1:1 | Accuracy | 0.991 | 1.0 | 0.989 | 1.0 | 0.984 | 0.999 | 0.977 | 0.994 | 0.957 | 0.975 |
|  | percentage | 0.153 | 0.009 | 0.256 | 0.047 | 0.403 | 0.149 | 0.584 | 0.344 | 0.809 | 0.638 |
| 3:2 | Accuracy | 0.988 | 1.0 | 0.981 | 1.0 | 0.976 | 0.996 | 0.966 | 0.99 | 0.943 | 0.964 |
|  | percentage | 0.123 | 0.007 | 0.21 | 0.039 | 0.339 | 0.122 | 0.497 | 0.29 | 0.73 | 0.569 |
| 2:1 | Accuracy | 0.961 | 1.0 | 0.961 | 0.991 | 0.96 | 0.99 | 0.943 | 0.979 | 0.917 | 0.938 |
|  | percentage | 0.081 | 0.005 | 0.146 | 0.027 | 0.242 | 0.086 | 0.388 | 0.221 | 0.618 | 0.475 |
| 5:2 | Accuracy | 0.919 | 1.0 | 0.93 | 0.987 | 0.924 | 0.981 | 0.911 | 0.952 | 0.876 | 0.894 |
|  | percentage | 0.055 | 0.004 | 0.099 | 0.017 | 0.17 | 0.059 | 0.289 | 0.16 | 0.516 | 0.397 |
| 3:1 | Accuracy | 0.861 | 0.833 | 0.876 | 0.949 | 0.867 | 0.959 | 0.87 | 0.921 | 0.824 | 0.84 |
|  | percentage | 0.037 | 0.001 | 0.068 | 0.01 | 0.121 | 0.039 | 0.219 | 0.12 | 0.433 | 0.342 |

## Discussion

We demonstrate that the proposed scCapsNet model performs well in the cell type prediction. The parallel fully connected neural networks could function like a feature extractor as convolutional neural networks in the original CapsNet model. Furthermore, the capsule part of our scCapsNet model forces the feature extraction part to capture local feature even specific to just one cell type. The model could provide the precise contribution of each extracted feature to the cell type recognition. Through the analysis of the internal weights of each neural network connected inputs and primary capsule, and with the information about the contribution of each extracted feature to the cell type recognition, the scCapsNet model could relate gene sets from inputs to cell types. Those type specific gene sets are vital for the cell type identity. The loss of those genes could significantly damage the model’s ability to recognize corresponding cell type.

To sum up, our scCapNet model could automatically extract features from data, and then use those extracted features to accomplish the cell type classification task and compute the exact contribution of each extracted feature to the type classification. Therefore, our scCapsNet model could potentially be used in the classification scenario where multiple information sources are available such as -omic datasets with data generated across different biological layers (e.g., transcriptomics, proteomics, metabolomics) [11]. By integrating the different information sources to classification, our scCapsNet model could provide the precise contribution of each information source. And it is seem that our scCapsNet model is more suitable for biological data than original CapsNet model.

The internal weights of the neural network that connect inputs to primary capsule is a matrix with rows represent genes. The real value vector of a row could be viewed as a low dimension embedding of a gene. The embedding of the same gene from different context that relates different primary capsule may vary substantially, due to the embedding is constrained by the function of the corresponding primary capsule. For example, if the corresponding primary capsule is responsible for the B cell recognition, then the genes associate with B cell identity would be embedded in similar way. Those embedding with specific purpose could be utilized in downstream analysis such as the research of gene function and gene-gene relationship.

In view of such high prediction accuracy to recognize mixed data, our scCapsNet model and the comparison model could potentially apply to wide range of research areas related to scRNA-seq. For example, cells develop or differentiate from stem cells to terminal differentiated cells through a trajectory. Along the trajectory, there are several states that have already been well characterized before. Training with those well defined states which could be viewed as landmarks, these two models could infer the locations of the testing cells on the trajectory by reporting two landmark states. Then, the testing cells could be viewed as at an intermediate state between the two reported landmarks. Like reporting the intermediate state of a cell, these two models could also be used to infer the intermediate position of a cell given several well defined landmark cells.

## Acknowledgments

This work was supported by grants from the National Key R&D Program of China [2018YFC0910402 to C.J.]; the National Natural Science Foundation of China [31571307 to C.J. and 61673070 to J.Z.]

## Supplementary Figures

**Figure S1:**
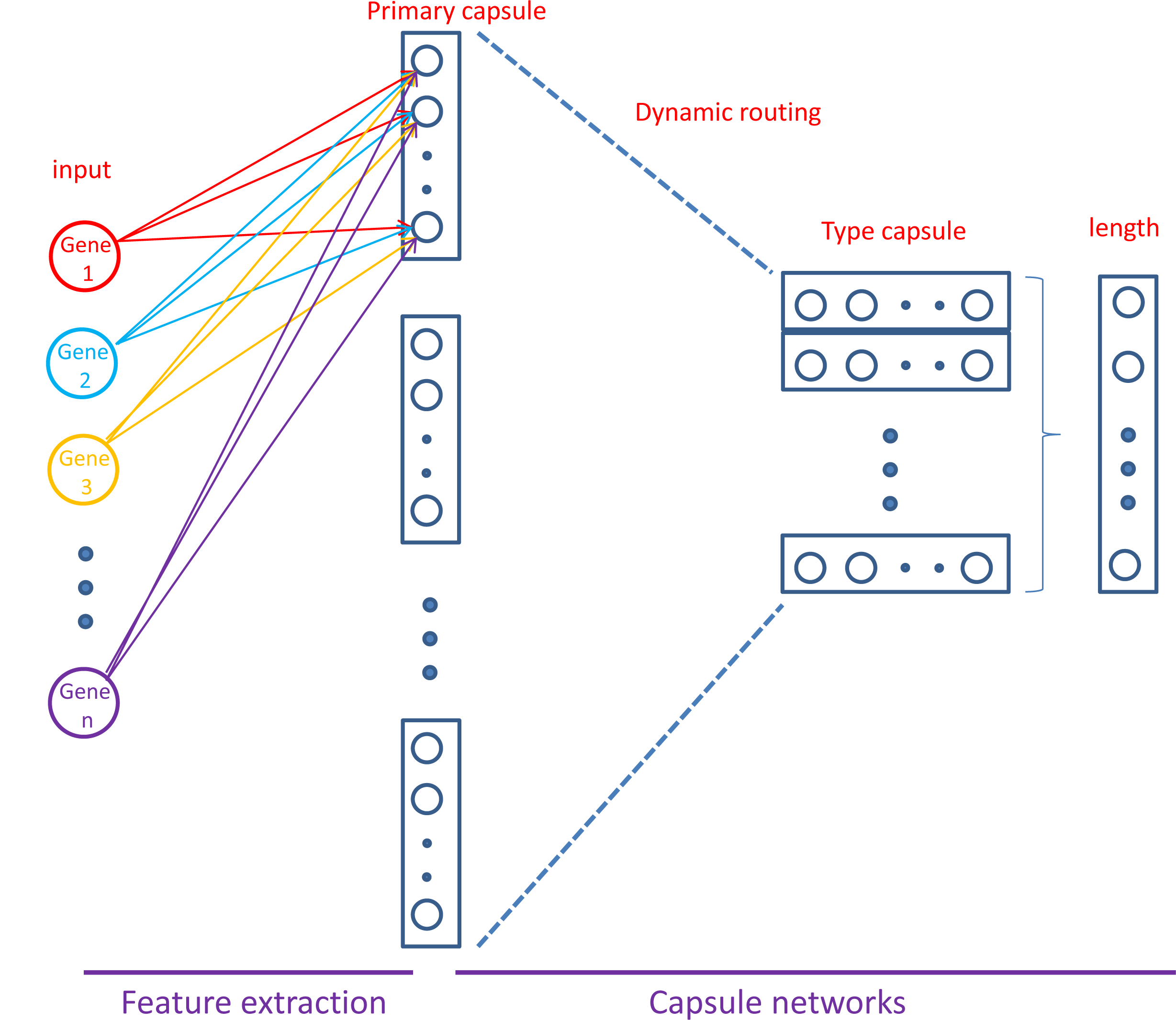
The visualization of internal weight of neural network which connect inputs and a primary capsule. Each gene could represent by weight related to it.

**Figure S2:**
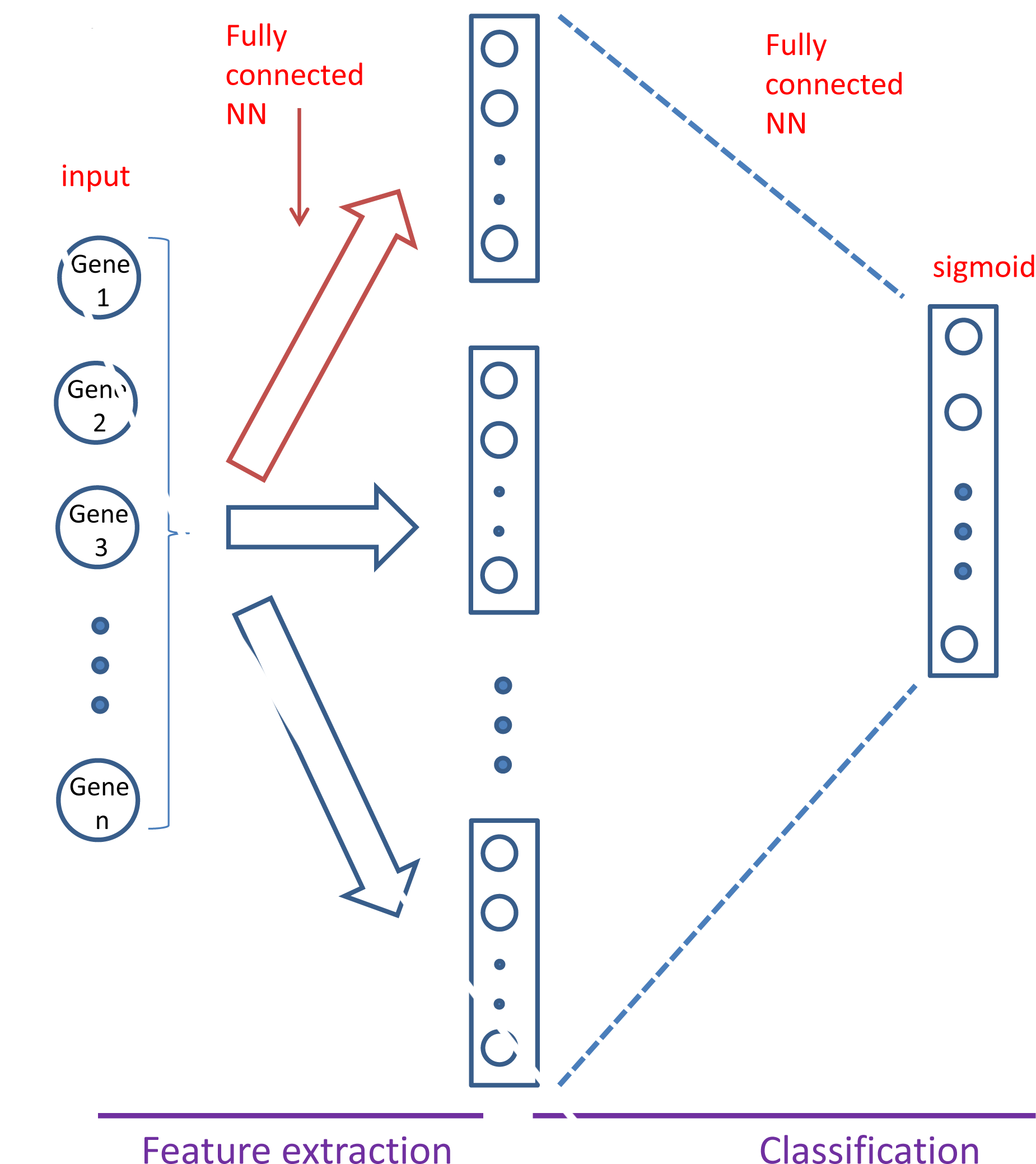
The architecture of the comparison neural network model. The comparison model retains the feature extraction layer of scCapsNet, but the capsule network part is substituted by a fully connected neural network with sigmoid activation function for classification.

**Figure S3:**
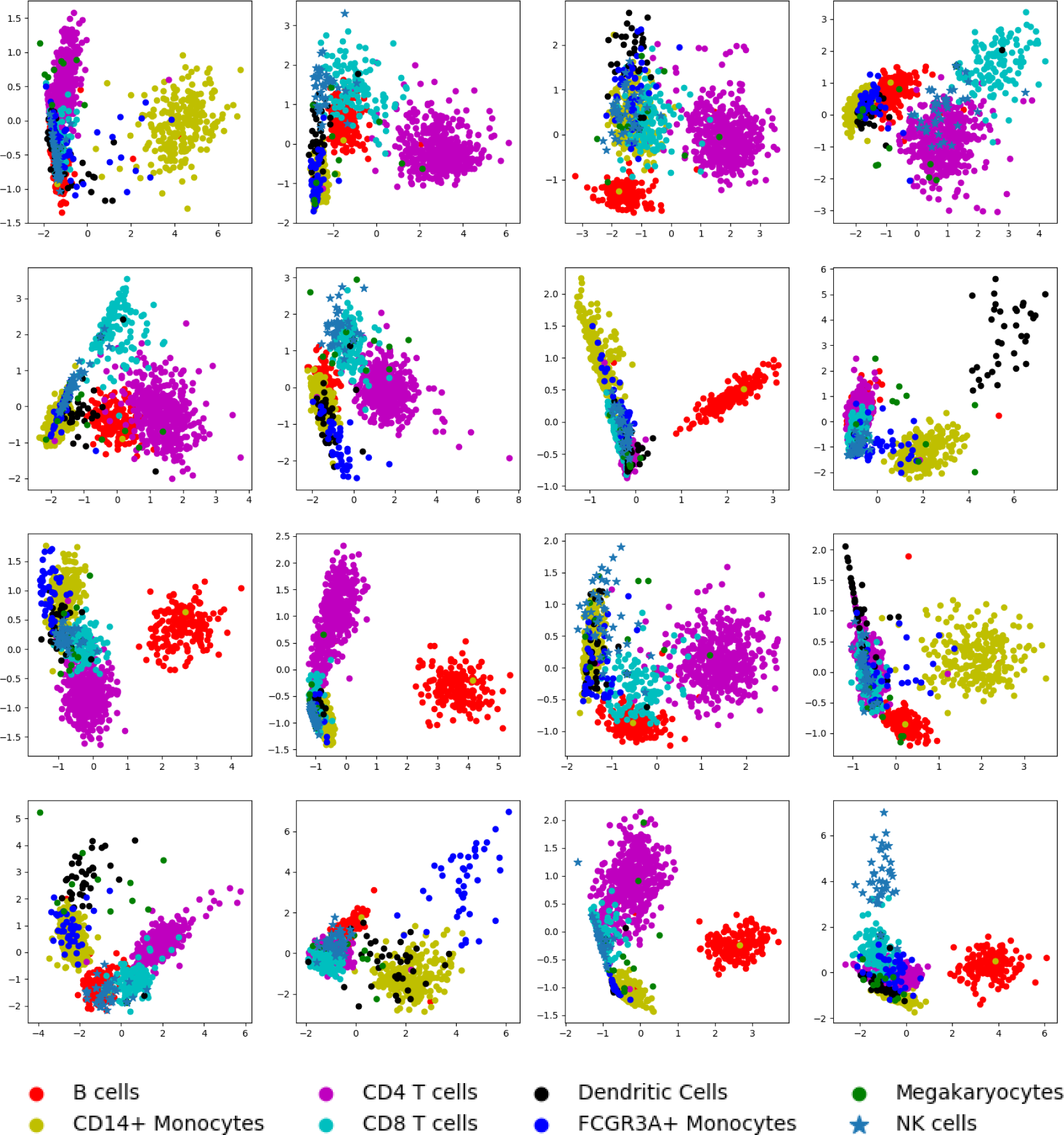
The 2D-PCA plots of primary capsules of input cells from validation set for PBMC dataset. Each subplot represents a primary capsule. The order of the subplot is from left to right and top to bottom with index from zero to fifteen.

**Figure S4:**
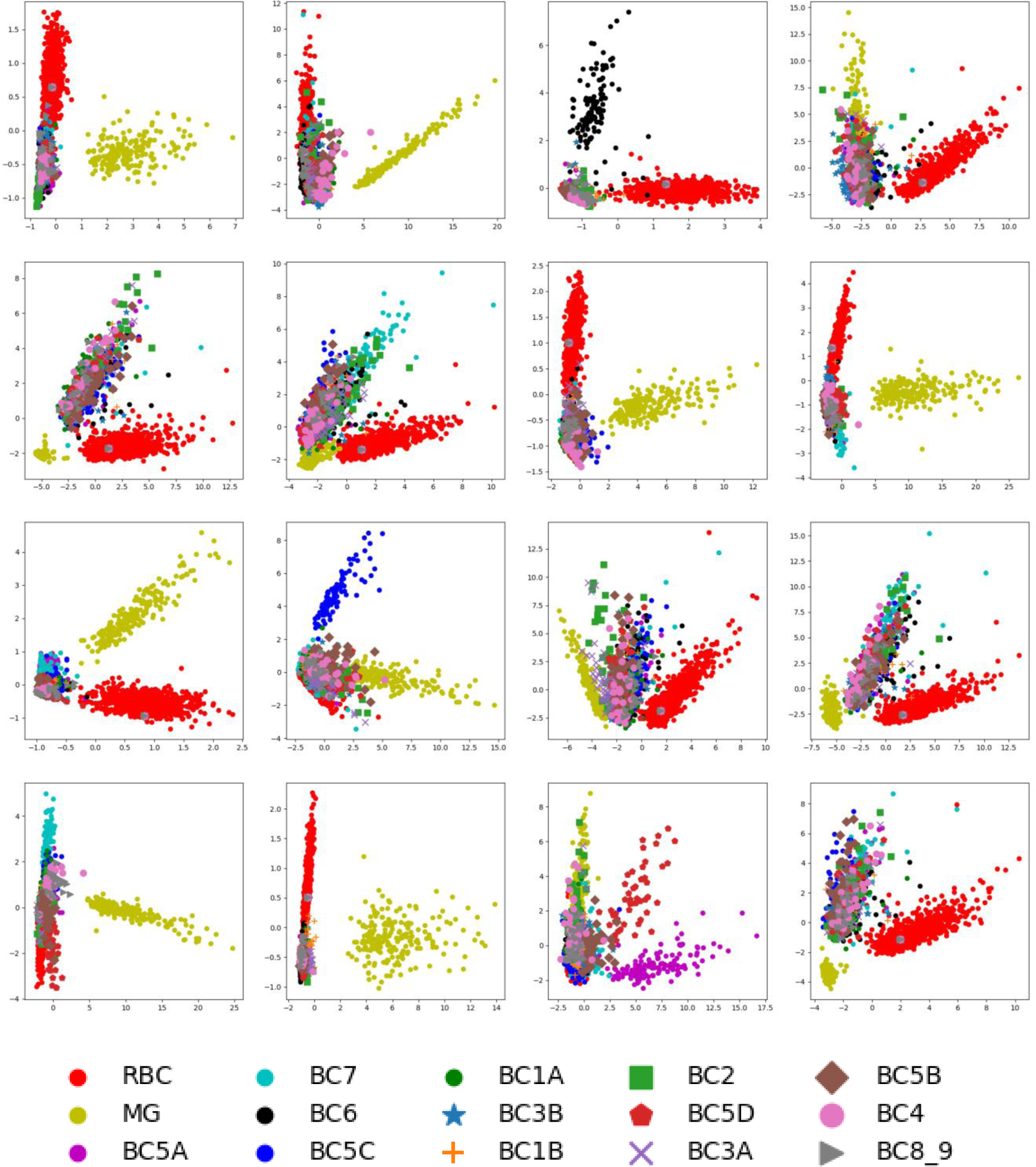
The 2D-PCA plots of primary capsules of input cells from validation set for mouse retina bipolar cells dataset. Each subplot represents a primary capsule. The order of the subplot is from left to right and top to bottom with index from zero to fifteen.

**Figure S5.**
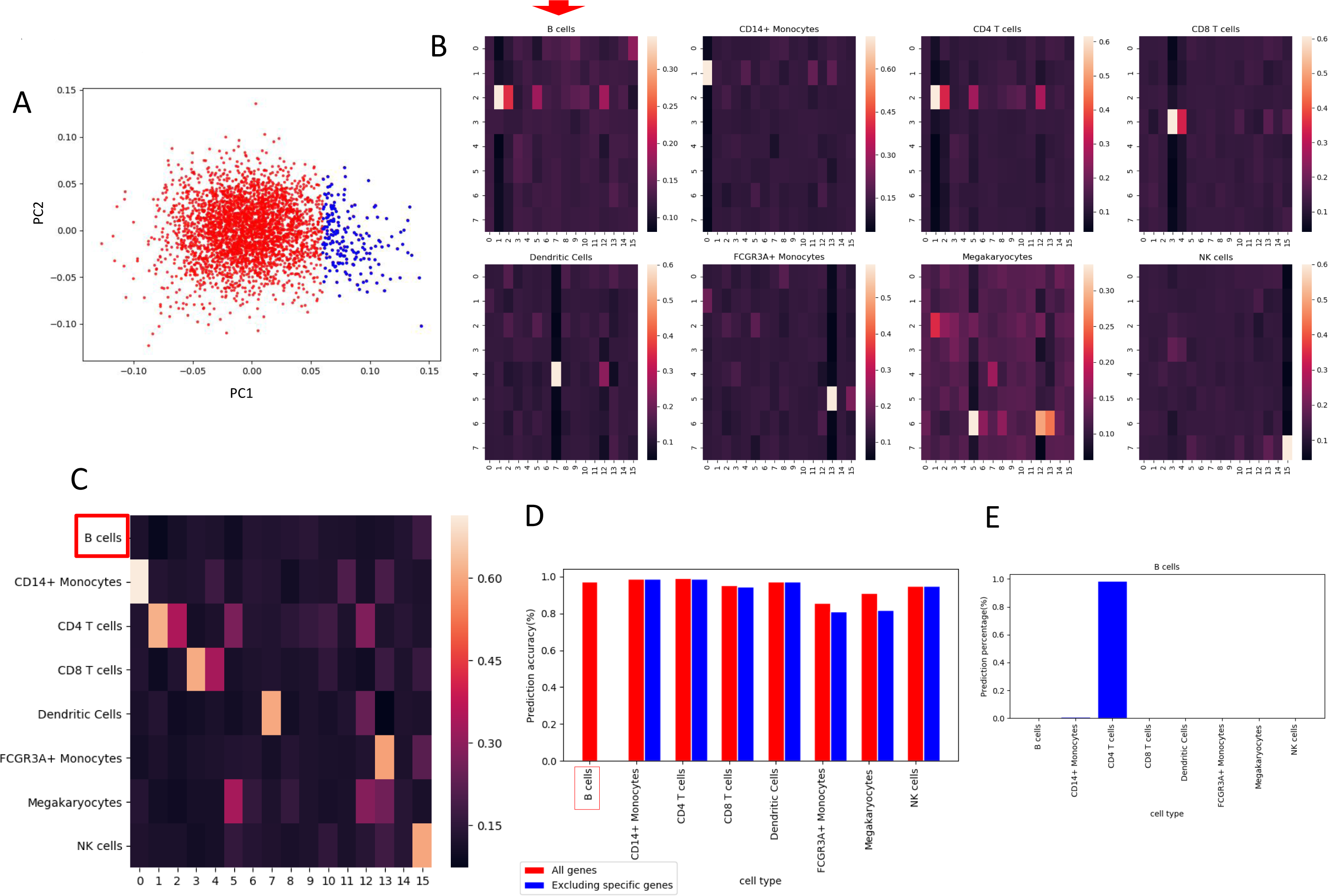
The identification of the gene sets specific for B cell, CD14+ monocytes, CD4 T cells, Dendritic cells, FCGR3A+ monocytes, and Megakaryocytes recognitions. **A**. The plot of the first two principle components (PC) of PCA which perform on internal weights of neural network that connect inputs and primary capsule nine, zero, one, seven, thirteen and five. Each dot represents a gene. Blue dots are chosen for genes exclusion. **B**. The heatmaps of coupling coefficients for deficit PBMC dataset in which the inputs value of blue dot genes are set to zero. **C**. The overall heatmap of average coupling coefficients with type capsules corresponding to a specific cell type input for deficit PBMC dataset. **D**. The comparison of prediction accuracy of each cell type between the untouched PBMC dataset and the deficit PBMC dataset. **E**. The proportion misclassified cell types of B cell, CD14+ monocytes, CD4 T cells, Dendritic cells, FCGR3A+ monocytes, and Megakaryocytes with deficit PBMC dataset.

**Figure S6.**
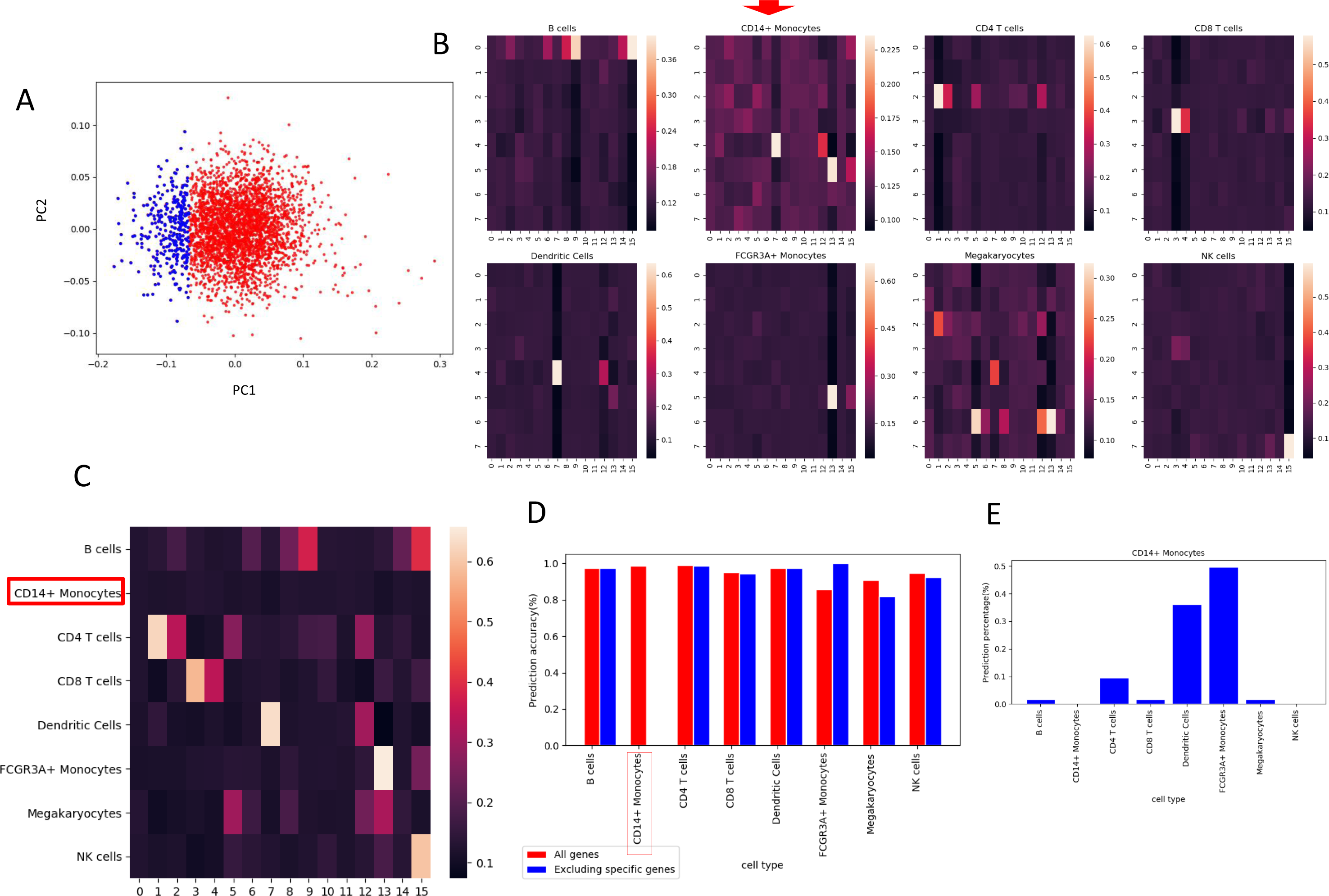
The identification of the gene sets specific for B cell, CD14+ monocytes, CD4 T cells, Dendritic cells, FCGR3A+ monocytes, and Megakaryocytes recognitions. **A**. The plot of the first two principle components (PC) of PCA which perform on internal weights of neural network that connect inputs and primary capsule nine, zero, one, seven, thirteen and five. Each dot represents a gene. Blue dots are chosen for genes exclusion. **B**. The heatmaps of coupling coefficients for deficit PBMC dataset in which the inputs value of blue dot genes are set to zero. **C**. The overall heatmap of average coupling coefficients with type capsules corresponding to a specific cell type input for deficit PBMC dataset. **D**. The comparison of prediction accuracy of each cell type between the untouched PBMC dataset and the deficit PBMC dataset. **E**. The proportion misclassified cell types of B cell, CD14+ monocytes, CD4 T cells, Dendritic cells, FCGR3A+ monocytes, and Megakaryocytes with deficit PBMC dataset.

**Figure S7.**
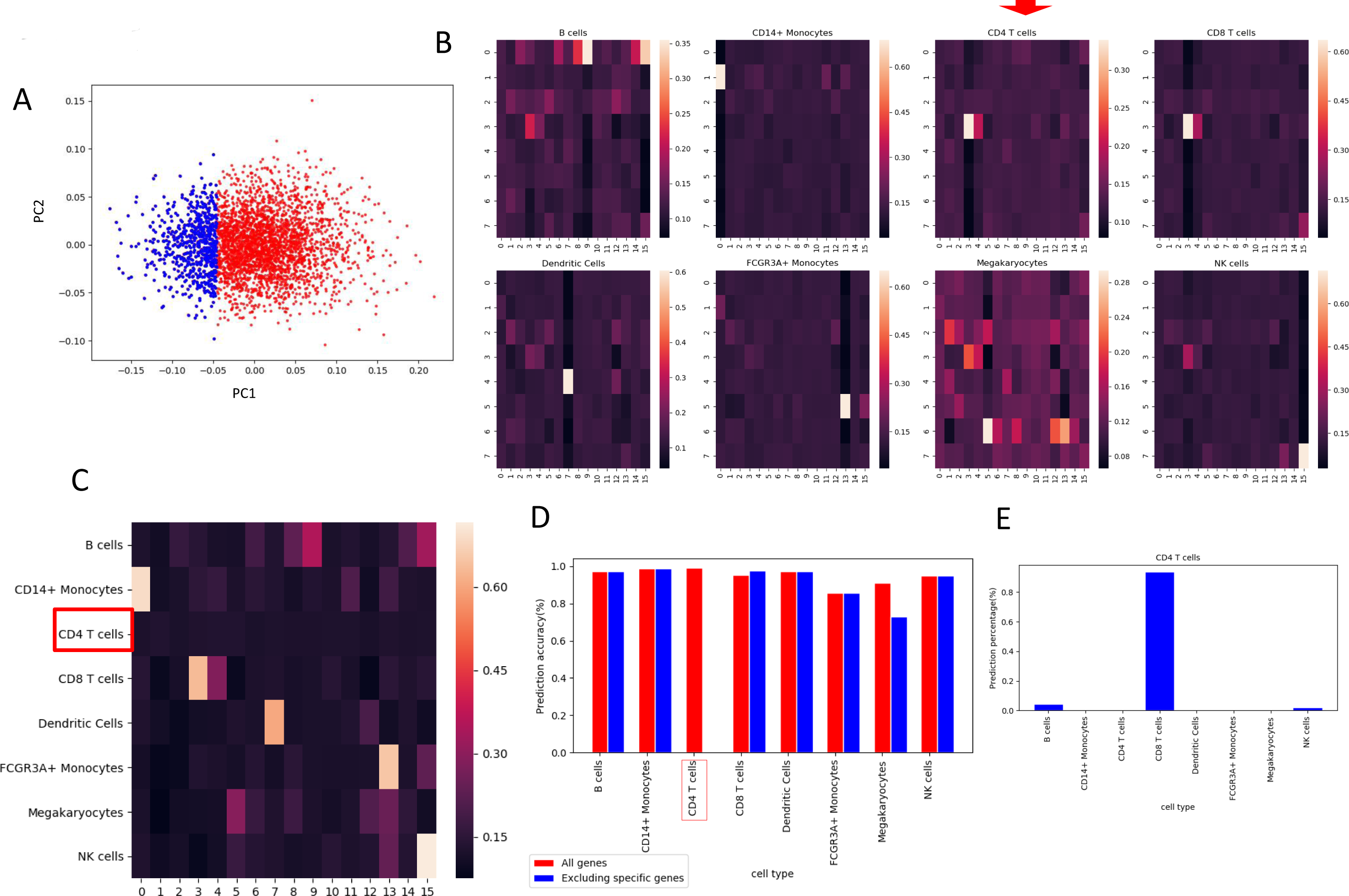
The identification of the gene sets specific for B cell, CD14+ monocytes, CD4 T cells, Dendritic cells, FCGR3A+ monocytes, and Megakaryocytes recognitions. **A**. The plot of the first two principle components (PC) of PCA which perform on internal weights of neural network that connect inputs and primary capsule nine, zero, one, seven, thirteen and five. Each dot represents a gene. Blue dots are chosen for genes exclusion. **B**. The heatmaps of coupling coefficients for deficit PBMC dataset in which the inputs value of blue dot genes are set to zero. **C**. The overall heatmap of average coupling coefficients with type capsules corresponding to a specific cell type input for deficit PBMC dataset. **D**. The comparison of prediction accuracy of each cell type between the untouched PBMC dataset and the deficit PBMC dataset. **E**. The proportion misclassified cell types of B cell, CD14+ monocytes, CD4 T cells, Dendritic cells, FCGR3A+ monocytes, and Megakaryocytes with deficit PBMC dataset.

**Figure S8.**
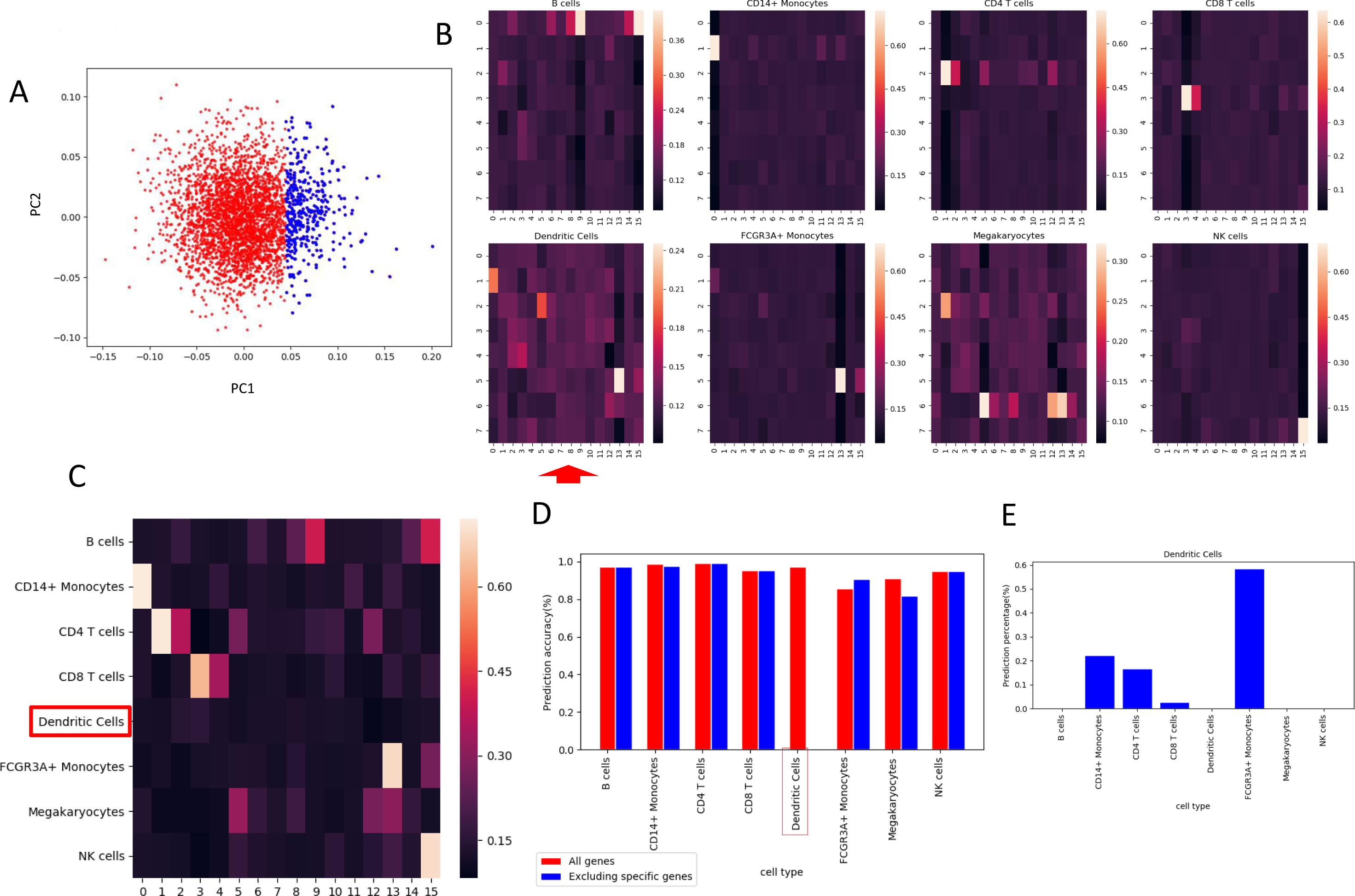
The identification of the gene sets specific for B cell, CD14+ monocytes, CD4 T cells, Dendritic cells, FCGR3A+ monocytes, and Megakaryocytes recognitions. **A**. The plot of the first two principle components (PC) of PCA which perform on internal weights of neural network that connect inputs and primary capsule nine, zero, one, seven, thirteen and five. Each dot represents a gene. Blue dots are chosen for genes exclusion. **B**. The heatmaps of coupling coefficients for deficit PBMC dataset in which the inputs value of blue dot genes are set to zero. **C**. The overall heatmap of average coupling coefficients with type capsules corresponding to a specific cell type input for deficit PBMC dataset. **D**. The comparison of prediction accuracy of each cell type between the untouched PBMC dataset and the deficit PBMC dataset. **E**. The proportion misclassified cell types of B cell, CD14+ monocytes, CD4 T cells, Dendritic cells, FCGR3A+ monocytes, and Megakaryocytes with deficit PBMC dataset.

**Figure S9.**
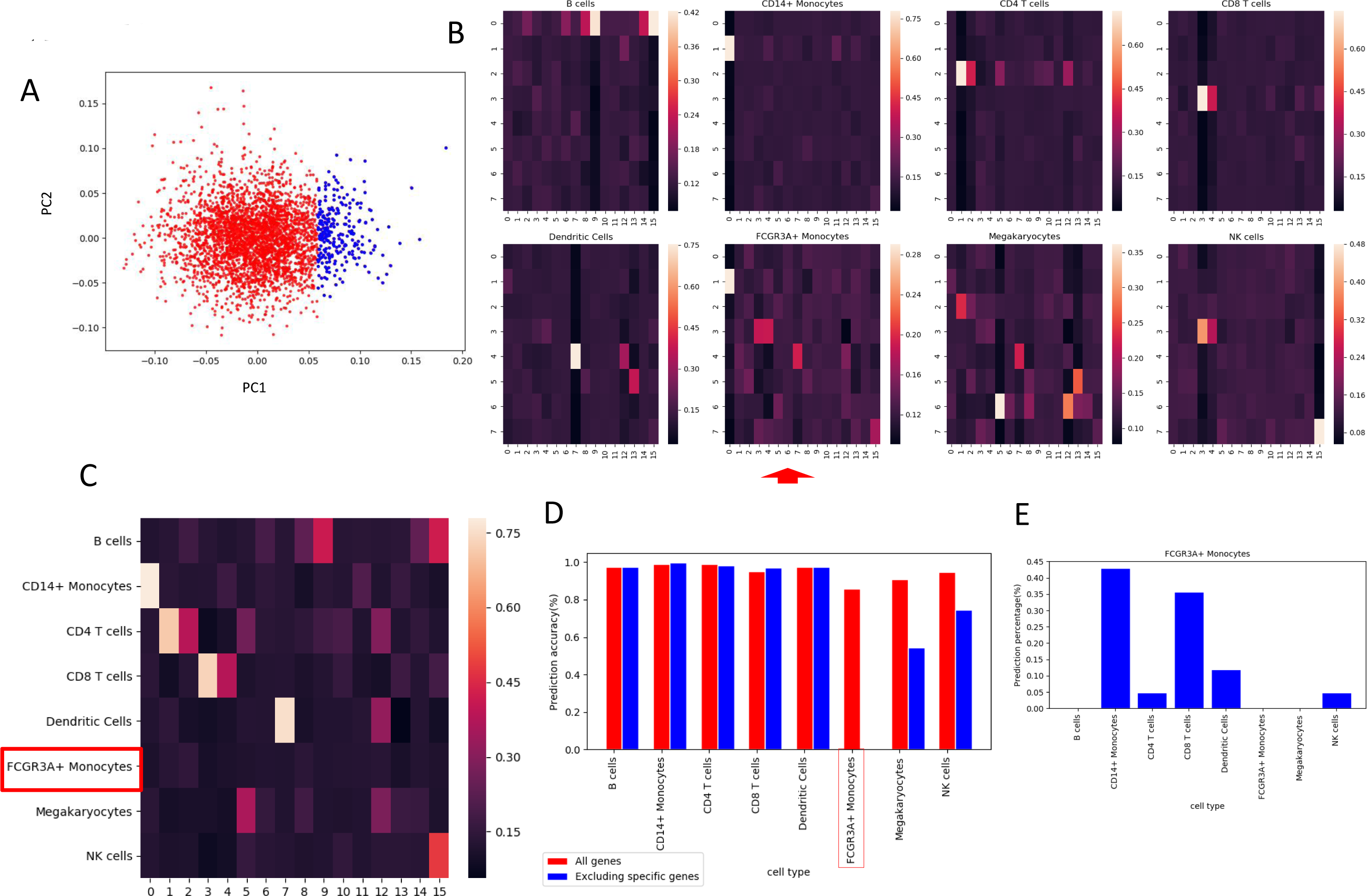
The identification of the gene sets specific for B cell, CD14+ monocytes, CD4 T cells, Dendritic cells, FCGR3A+ monocytes, and Megakaryocytes recognitions. **A**. The plot of the first two principle components (PC) of PCA which perform on internal weights of neural network that connect inputs and primary capsule nine, zero, one, seven, thirteen and five. Each dot represents a gene. Blue dots are chosen for genes exclusion. **B**. The heatmaps of coupling coefficients for deficit PBMC dataset in which the inputs value of blue dot genes are set to zero. **C**. The overall heatmap of average coupling coefficients with type capsules corresponding to a specific cell type input for deficit PBMC dataset. **D**. The comparison of prediction accuracy of each cell type between the untouched PBMC dataset and the deficit PBMC dataset. **E**. The proportion misclassified cell types of B cell, CD14+ monocytes, CD4 T cells, Dendritic cells, FCGR3A+ monocytes, and Megakaryocytes with deficit PBMC dataset.

**Figure S10.**
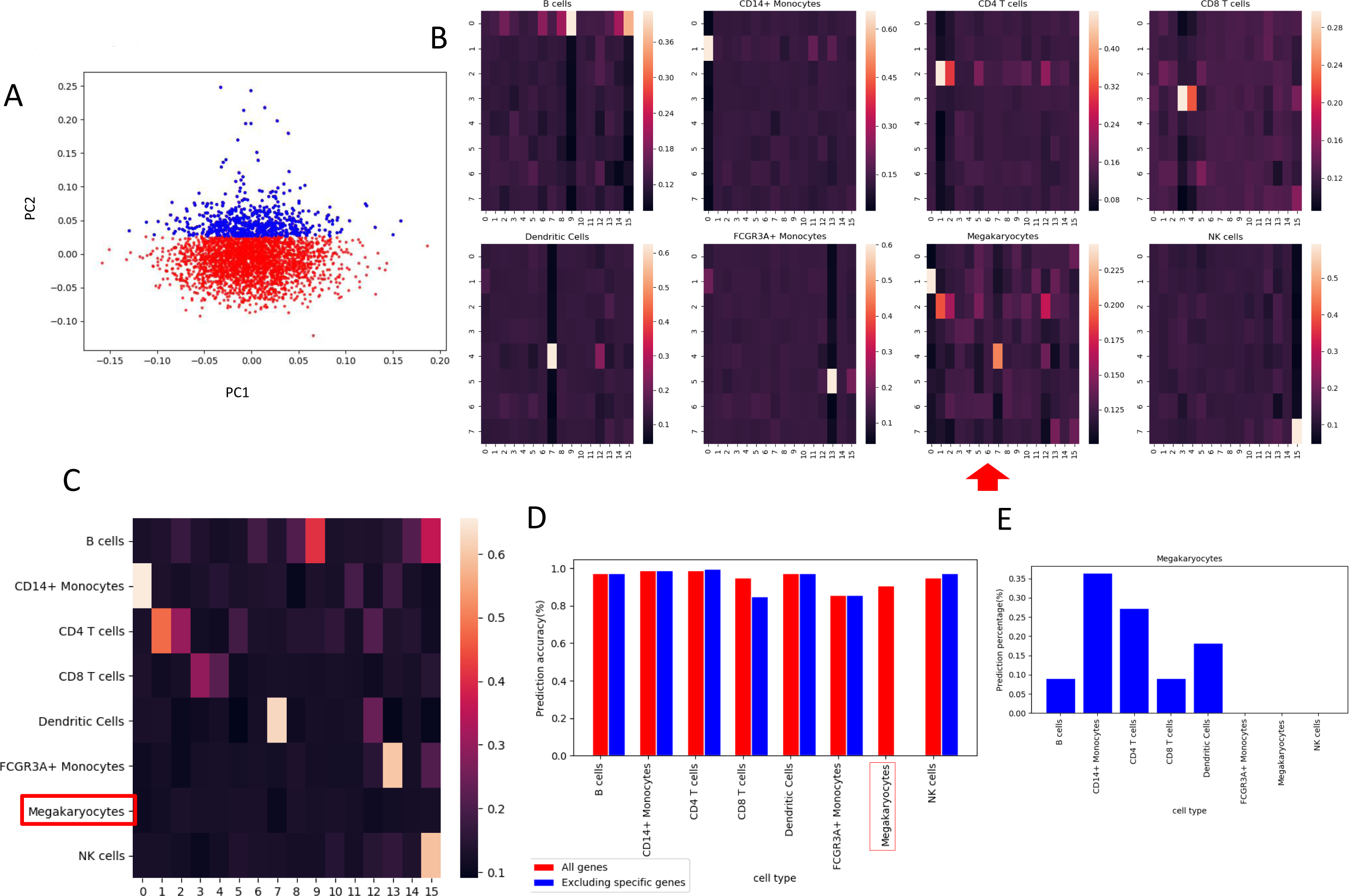
The identification of the gene sets specific for B cell, CD14+ monocytes, CD4 T cells, Dendritic cells, FCGR3A+ monocytes, and Megakaryocytes recognitions. **A**. The plot of the first two principle components (PC) of PCA which perform on internal weights of neural network that connect inputs and primary capsule nine, zero, one, seven, thirteen and five. Each dot represents a gene. Blue dots are chosen for genes exclusion. **B**. The heatmaps of coupling coefficients for deficit PBMC dataset in which the inputs value of blue dot genes are set to zero. **C**. The overall heatmap of average coupling coefficients with type capsules corresponding to a specific cell type input for deficit PBMC dataset. **D**. The comparison of prediction accuracy of each cell type between the untouched PBMC dataset and the deficit PBMC dataset. **E**. The proportion misclassified cell types of B cell, CD14+ monocytes, CD4 T cells, Dendritic cells, FCGR3A+ monocytes, and Megakaryocytes with deficit PBMC dataset.

**Figure S11:**
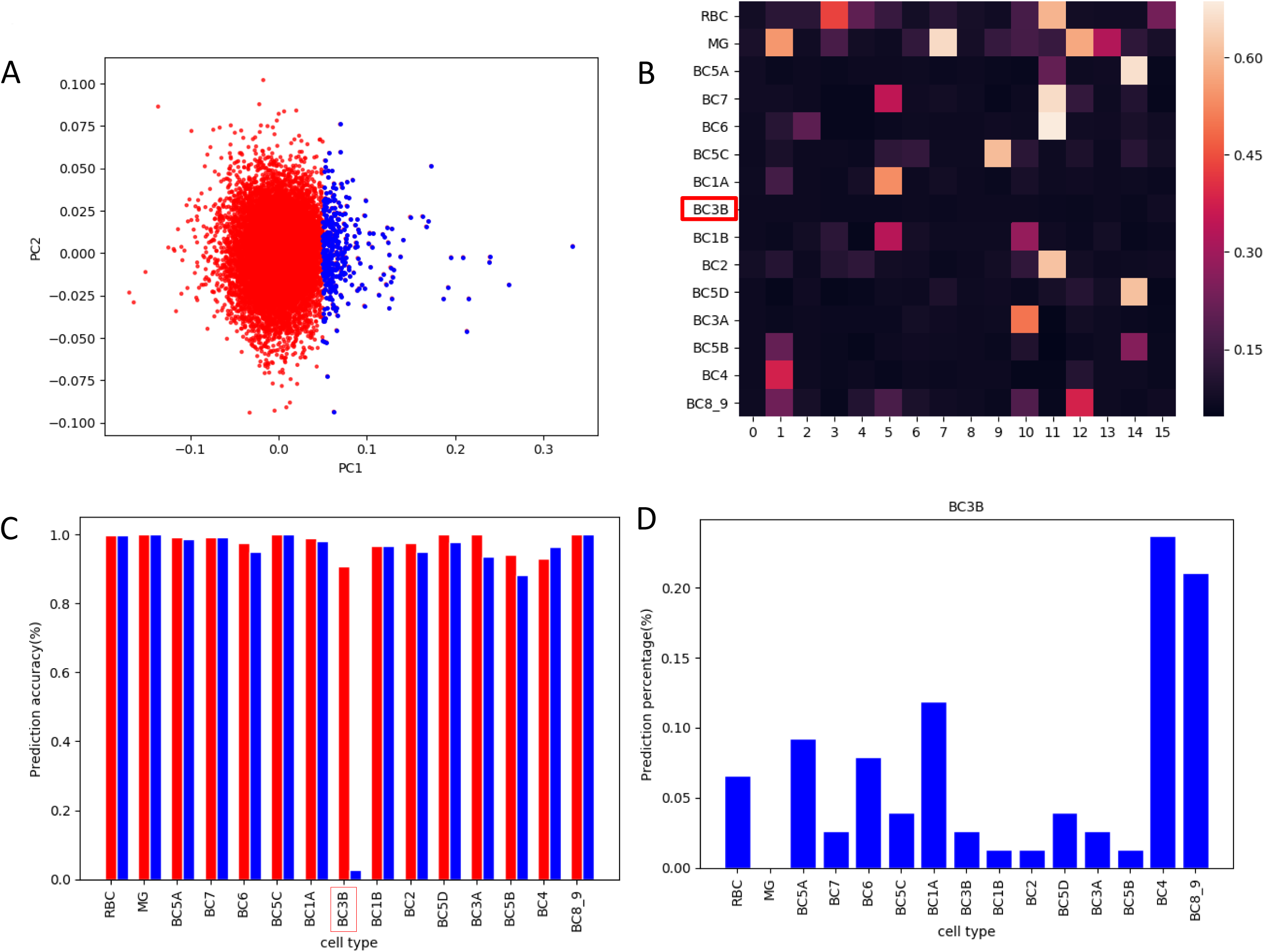
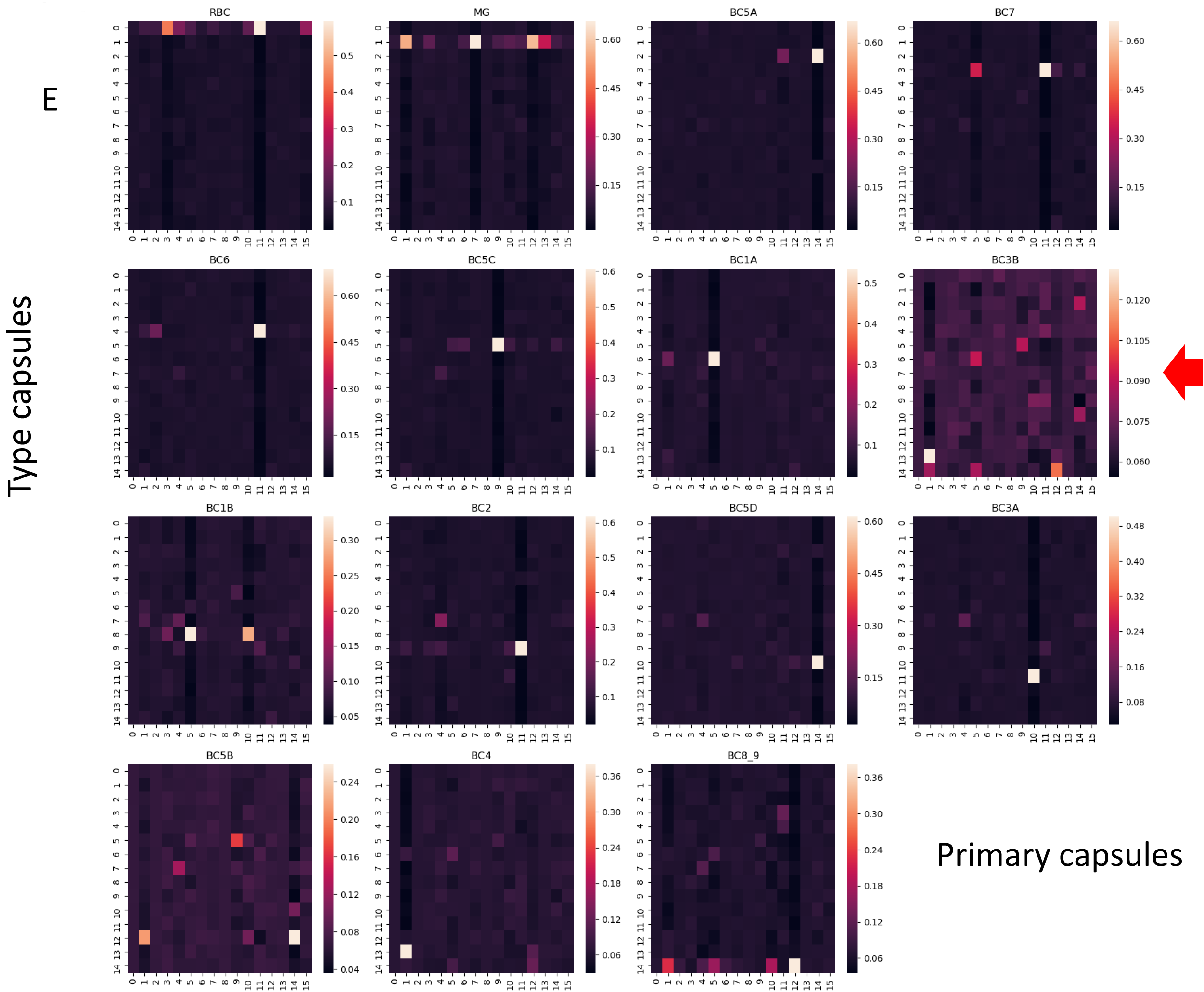
The identification of the gene set specific for BC3B cells, recognitions. **A**. The plot of the first two principle components (PC) of PCA which perform on internal weights of neural network that connect inputs and primary capsule. Each dot represents a gene. Blue dots are chosen for genes exclusion. **B**. The overall heatmap of average coupling coefficients with type capsules corresponding to a specific cell type input for deficit mouse retinal bipolar neurons dataset. **C**. The comparison of prediction accuracy of each cell type between the untouched mouse retinal bipolar neurons dataset and the deficit mouse retinal bipolar neurons dataset. **D**. The proportion misclassified cell types BC3B with deficit mouse retinal bipolar neurons dataset. **E.** The heatmaps of coupling coefficients for deficit mouse retinal bipolar neurons dataset in which the inputs value of blue dot genes are set to zero.

**Figure S12.**
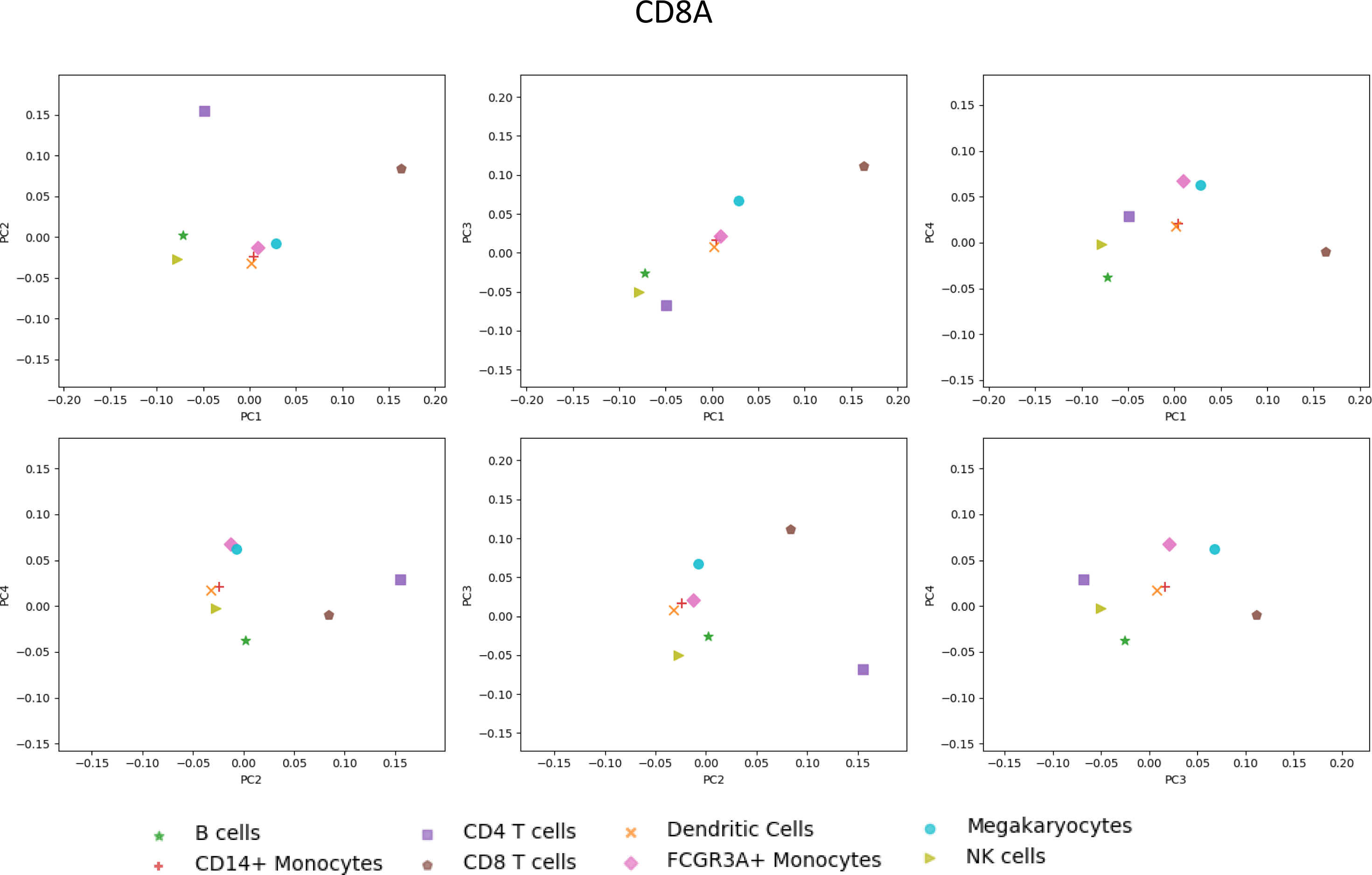
The position of CD8A, CD14 and CD19 on The plots of different pairs of principle components of Figure 8B.

**Figure S13.**
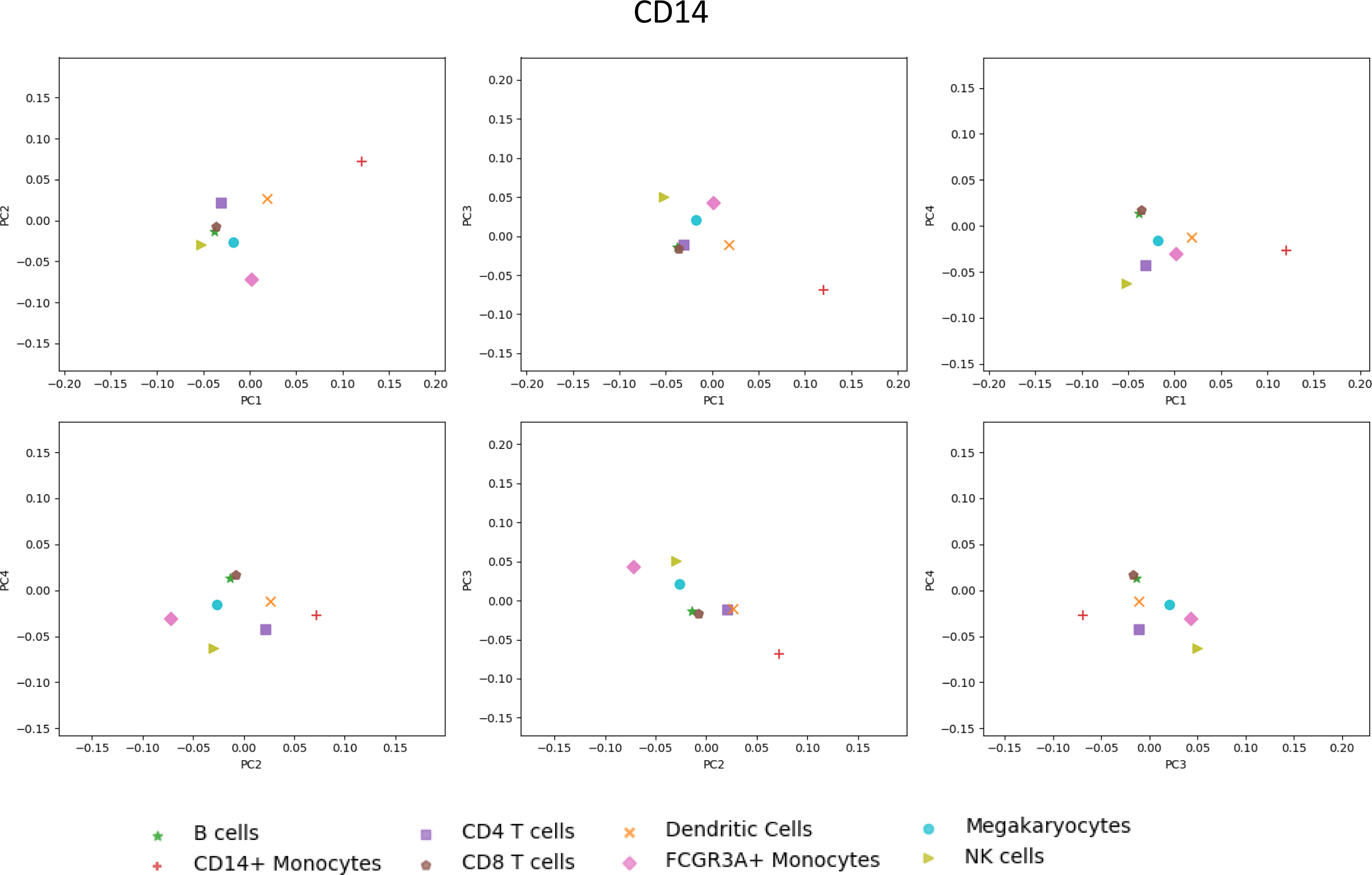
The position of CD8A, CD14 and CD19 on The plots of different pairs of principle components of Figure 8B.

**Figure S14.**
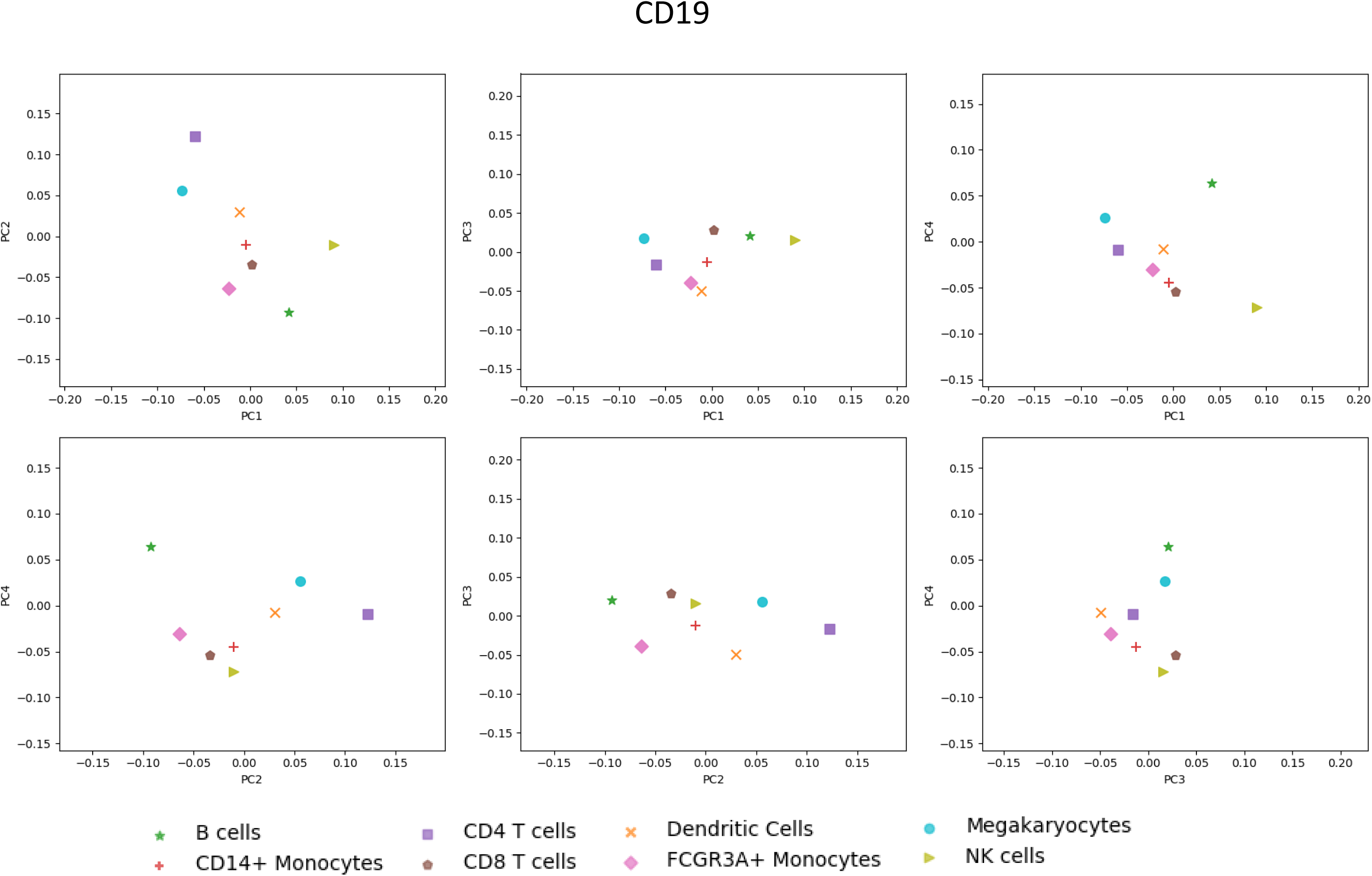
The position of CD8A, CD14 and CD19 on The plots of different pairs of principle components of Figure 8B.

